# Actin-dependent mechanotransduction controls nucleocytoplasmic partitioning of DNMT3a through ERK1/2 signaling during cutaneous wound healing

**DOI:** 10.64898/2026.01.15.699481

**Authors:** Johan Ajnabi, Binita Dam, Ekta Gupta, Tirthankar Saha, Abhik Dutta, Saurav Kumar, Ashish Gupta, Dasaradhi Palakodeti, Colin Jamora

## Abstract

Cutaneous wound healing is a multifaceted physiological process that requires a cell state transition from homeostasis to tissue repair. An important contributor to this reprogramming is the wound-induced mechanical cues that are perceived by epidermal keratinocytes. Previously we identified that the nuclear translocation of the *de novo* DNA methyltransferase 3A (DNMT3a) upon wounding as an important regulator of this cell-state transition. However, the molecular mechanisms linking mechanotransduction to epigenetic regulation remain incompletely understood. Here we show that under homeostasis, active ERK1/2 phosphorylates DNMT3a, intramolecularly masking its nuclear localization signal (NLS), resulting in its cytoplasmic sequestration. Upon wounding actin cytoskeleton remodeling leads to the downregulation of the Extracellular Signal-Related Kinase 1/2 (ERK 1/2) pathway. Wound-induced ERK1/2 inactivation unmasks the NLS, enabling DNMT3a nuclear translocation. Collectively, these findings define a mechanotransduction-driven signaling axis linking cytoskeletal dynamics to epigenetic regulation and confirms an active role for differentiated keratinocytes in initiating early wound repair programs.

## Introduction

Tissue repair in multicellular organisms is a product of an exquisite coordination of mechanical cues, biochemical signaling, and transcriptional networks. A growing body of evidence indicates that epigenetic mechanisms – including DNA methylation, histone modifications and non-coding RNAs – serve as the molecular interface that encodes and stabilizes these transcriptional programs during wound healing and regeneration.^1–3^ For instance, DNA methyltransferases DNMT1 and DNMT3B are upregulated during corneal epithelial wound healing, leading to a global DNA hypermethylation.^4^ In pathological contexts, such as diabetes, impaired wound healing is associated with DNMT1 upregulation and hypermethylation-driven dysregulation of Notch1, PU.1, Klf4, and TLR2, genes critical for stem cell function, immune responses, and tissue regeneration.^5^ Abnormal activity of histone-modifying enzymes – including Jmjd3, MLL1, Polycomb group proteins, SETD2, and HDACs – has been shown to regulate macrophage and keratinocyte growth and function, thereby influencing distinct stages of the wound-healing response.^6–11^ Non-coding RNAs, particularly microRNAs, further modulate wound repair by fine-tuning the expression of epigenetic regulators and downstream transcriptional networks in inflammatory and proliferative phases of wound healing.^12,13^ Collectively, these findings illustrate that epigenetic modifications are central coordinators of effective wound healing. Despite extensive evidence linking epigenetic regulation to wound healing, how mechanical forces generated during tissue injury dynamically control epigenetic regulators remains an unanswered fundamental question. Although wounding induces profound changes in cellular tension, cell shape, and cytoskeletal organization, the molecular mechanisms that connect these physical cues to chromatin-modifying enzymes are poorly defined.

The epidermis provides an ideal system to investigate the link between mechanotransduction and epigenetic regulation. Under homeostatic conditions, differentiated keratinocytes form a mechanically cohesive epithelial sheet, supported by adherens junctions and a highly organized actin cytoskeleton that sustains elevated intracellular tension.^14,15^ Upon wounding, keratinocytes undergo rapid transcriptional reprogramming that enables them to transit from a terminally differentiated state to a reparative, migratory phenotype.^16–18^

We previously identified DNMT3a, a *de novo* DNA methyltransferase, as a critical modulator of the transcriptome landscape required for the initiation of the tissue repair process.^17^ Notably, in differentiated keratinocytes DNMT3a is predominantly localized in the cytoplasm under homeostatic conditions but rapidly translocates to the nucleus following wounding. The nuclear translocation of DNMT3a was sufficient to modulate known wound healing genes and caused partial dedifferentiation of keratinocytes that is consistent with a tissue repair program.^17^ Given DNMT3a’s role in establishing *de novo* methylation patterns during differentiation and stress responses, this observation suggested that its epigenetic activity may be dynamically regulated at the level of subcellular localization.^19,20^ These findings raised the intriguing possibility that the nuclear availability of DNMT3a is controlled by the mechanical state of the cell, directly linking physical cues to epigenetic reprogramming. However, the upstream signals governing DNMT3a localization and activation in response to injury remained unknown.

## Results

### Wound-induced actin remodelling is necessary for nuclear localization of DNMT3a

We have previously reported that wound-induced cellular tension release leads to nuclear localization of DNMT3a in keratinocytes.^17^ This localization is observed at the wound edge within the differentiated layers of the epidermis.^17^ The molecular mechanism governing this mechanoregulation, however, remains unclear. In order to determine whether DNMT3a acts as an immediate effector of mechanical tension release or as a downstream target of secondary signaling cascades, a temporal resolution is required. To address this, we performed a kinetic analysis of DNMT3a nuclear translocation following wounding. On a layer of differentiated keratinocytes, we performed an *in vitro* scratch wound assay and observed the nucleo-cytoplasmic localization of DNMT3a at 0-, 2-, 5-, 10- and 15-minutes post wounding (Figure 1A). The ratio of nuclear and cytoplasmic integrated density of DNMT3a is quantified and represented as DNMT3a (nuc/cyt) ratio (Figure 1B).^21^ A calculated DNMT3a (nuc/cyt) ratio of 1 (indicated by the dashed line) defines the threshold for subcellular localization, with values below and above this threshold corresponding to predominant cytoplasmic and nuclear localization, respectively. Scratch wounding induced nuclear localization of DNMT3a in cells within 200 µm of the wound edge, detectable at 5 minutes, increasing at 10- and 15-minutes post wounding (Figure 1A and 1B). No changes were observed in distal cells (>1000 µm from the wound edge) across the time course (Figure 1B and S1A). Spatiotemporal analysis also revealed a gradient of DNMT3a nuclear localization from the wound edge, with maximal nuclear accumulation within 200 µm (Figure S1B and S1C). Localization progressively decreased with distance, and beyond 600 µm DNMT3a was predominantly cytoplasmic (Figure S1B and S1C). These data indicate that DNMT3a nuclear translocation is spatially and temporally regulated.

**Figure 1:**
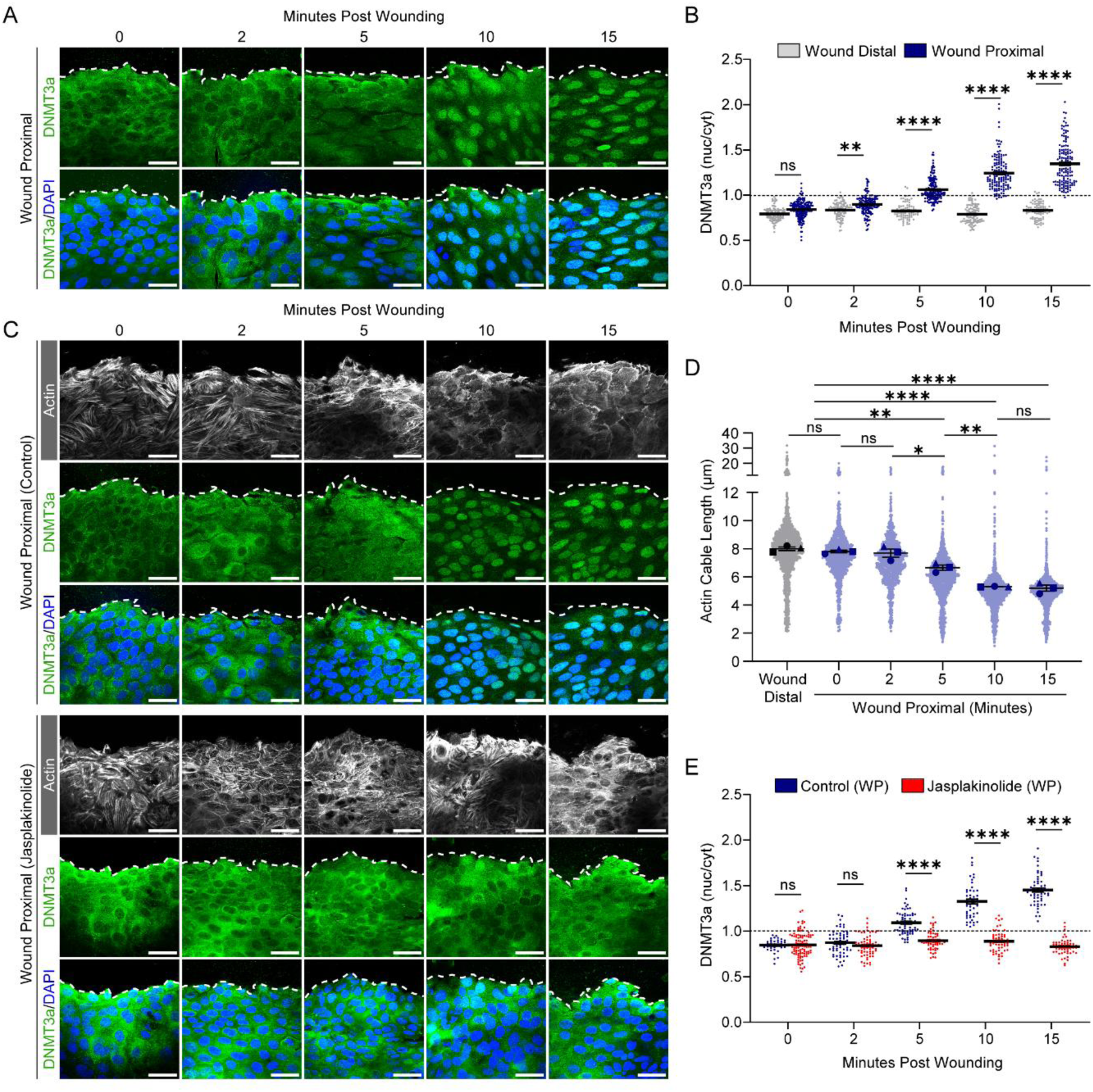
Actin remodleing induced by wounding is required for nuclear localization of DNMT3a. **(A)** DNMT3a immunostaining (green) and DAPI (blue) at wound proximal (WP) site (≤ 200 µm from wound edge. **(B)** Quantification and kinetics of DNMT3a subcellular localization represented as DNMT3a (nuc/cyt) ratio. Dashed line at Y=1 defines the threshold for subcellular localization. **(C)** Top panel: phalloidin staining (grey) shows time-dependent actin cytoskeleton remodelling correlated with nuclear localization of DNMT3a (green); Bottom panel: similar immunostaining with Jasplakinolide treatment prior wounding **(D)** Quantification of actin cable length upon wounding. **(E)** Quantification of DNMT3a (nuc/cyt) ratio upon wounding with or without Jasplakinolide treatment. Scale bars: 50 µm for all images. The data are represented as mean ± s.e.m. n=6 for **(A)**, **(B)** and **(D)**, n=3 for **(C)** and **(E)**. P-values in **(B)**, **(D)** and **(E)** were calculated using 2-way ANOVA with Sidak’s multiple comparison test where ns= P > 0.05, *= P ≤ 0.05, **= P ≤ 0.01, ***= P ≤ 0.001, ****= P ≤ 0.000

We next investigated how wound-induced mechanical cues regulate the spatiotemporal localization of DNMT3a. Differentiated keratinocytes form an epithelial sheet with strong junctional adhesion and interconnected actin cables, which maintain high cellular tension.^14,15,22^ Upon wounding, these actin cables at the wound edge are rapidly dissolved, releasing cellular tension.^23,24^ Consistent with this, we observed swift actin remodeling in cells at the wound edge (Figure 1C). Quantitative analysis revealed a significant reduction in mean actin cable length at the proximal wound site (≤200 µm) within 5 minutes of wounding, which progressively declined and plateaued by 10 minutes (Figure 1D). Whereas it remained largely unchanged over time in the wound distal region (Figure S1F). The kinetics of the decrease in filamentous actin length temporally overlapped with the nuclear accumulation of DNMT3a (Figure 1B, 1C and 1D). Linear regression analysis confirmed a strong correlation between actin cable shortening and DNMT3a nuclear localization in a time-dependent manner (Figure S1E). Similar correlation was observed in the spatially resolved image showing a gradient of nuclear localization from the wound edge that correlates inversely with the actin cable length shortening (Figure S1B, S1C and S1D). The spatial gradient of actin remodelling and DNMT3a nuclear localization from the wound edge closely parallels the gradient of tension release observed in migrating MDCK cell sheets following wounding.^25^ Collectively, these findings suggest that DNMT3a nuclear translocation is coupled to wound-induced actin remodeling.

To determine whether wound-induced actin remodeling is required for DNMT3a nuclear localization, cells were treated with 100 nM Jasplakinolide, an actin-stabilizing compound,^26^ prior to wounding. Jasplakinolide treatment prevented wound-induced actin remodeling (Figure S1G) and concomitantly inhibited DNMT3a nuclear accumulation (Figure 1C and 1E). These results indicate that dynamic actin remodeling is necessary for the nuclear localization of DNMT3a following wounding.

### Wound-induced actin remodeling is sufficient for nuclear localization of DNMT3a

Given that Jasplakinolide-mediated stabilization of the actin cytoskeleton prevents wound-induced DNMT3a nuclear accumulation, we next examined whether actin disruption alone is sufficient to drive this process. Treatment with Latrunculin A, which sequesters actin monomers and blocks polymerization,^27^ induced nuclear translocation of DNMT3a (Figure 2A and 2B). Similarly, Cytochalasin D, which caps the barbed (+) end of actin filaments to inhibit polymerization,^28^ also triggered DNMT3a nuclear localization (Figure S2A and S2B). These results demonstrate that actin disruption, independent of other wound-associated cues, is sufficient for DNMT3a nuclear translocation.

**Figure 2:**
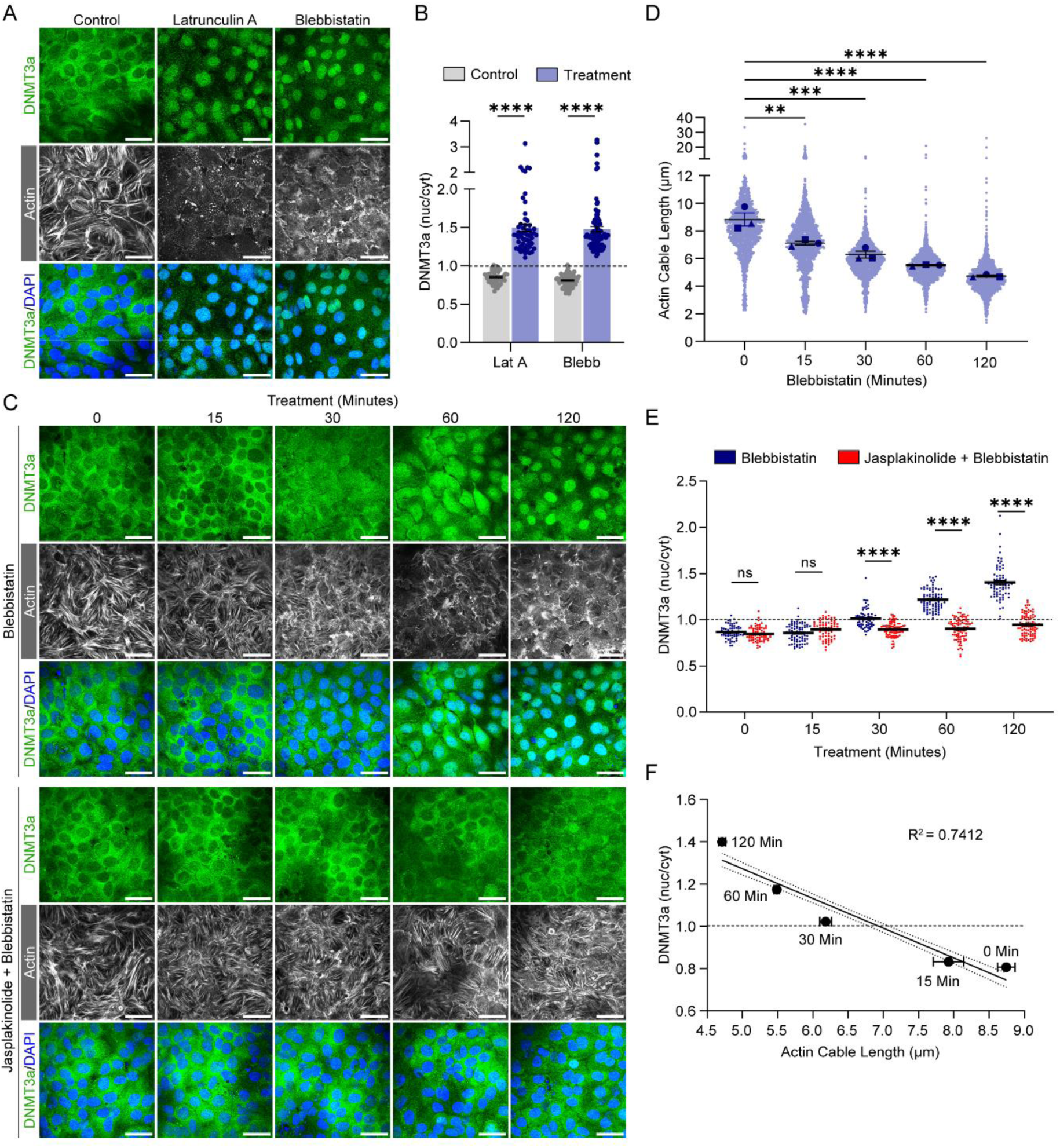
Partial actin disruption is sufficient to drive nuclear localization of DNMT3a. **(A)** DNMT3a immunostaining (green) with phalloidin (grey), and DAPI (blue) upon Latrunculin A and Blebbistatin treatment. **(B)** Quantification of DNMT3a (nuc/cyt) upon Latrunculin A and Blebbistatin treatment. **(C)** DNMT3a immunostaining (green) with actin (grey) and DAPI (blue) upon Blebbistatin treatment (top panel) or Jasplakinolide + Blebbistatin treatment (bottom panel). **(D)** Quantification of actin cable length upon Blebbistatin treatment. **(E)** Quantification and kinetics of DNMT3a subcellular localization represented as DNMT3a (nuc/cyt) ratio upon Blebbistatin (blue) and Jasplakinolide + Blebbistatin (red) treatment. **(F)** Linear regression analysis between actin cable length and nuclear localization of DNMT3a upon Blebbistatin treatment. Scale bar: 50 µm for all images. The data are represented as mean ± s.e.m. n=4. P-values in **(B)** calculated using unpaired Welch’s t test, in **(D)** and **(E)** were calculated using two-way ANOVA with Tukey’s multiple comparison test where ns= P > 0.05, *= P ≤ 0.05, **= P ≤ 0.01, ***= P ≤ 0.001, ****= P ≤ 0.0001.

Because scratch wounding does not completely disrupt the actin cytoskeleton (Figure 1C and 1D), we investigated whether partial actin remodeling could likewise stimulate DNMT3a nuclear localization. Treatment with Blebbistatin, a myosin II inhibitor that disassembles actin bundles and mimics wound-like shortening of actin cables (Figure 2C, 2D and S2C), led to DNMT3a nuclear accumulation (Figure 2A and 2B). Partial actin dissolution induced by the ROCK inhibitor Y-27632 produced a similar outcome (Figure S2A and S2B). These findings indicate that complete actin disruption is not required. Instead, partial remodeling of actin bundles is sufficient to induce DNMT3a nuclear translocation.

Since the wound response is highly localized (within a few hundred micrometres of the wound edge), signal-to-noise limitations precluded biochemical assays. To address this, we compared actin remodeling induced by various destabilizing agents and identified Blebbistatin as most closely mimicking wound-induced actin shortening (Figure S2C). We therefore employed Blebbistatin as a wound proxy for subsequent biochemical analyses. Time-course experiments revealed that Blebbistatin treatment, similar to proximal wounding, caused progressive shortening of actin cables (Figure 2C and 2D) accompanied by concomitant nuclear accumulation of DNMT3a (Figure 2C and 2E). Importantly, this reflected a redistribution rather than altered protein abundance, as DNMT3a levels remained constant throughout the time course (Figure S2D). Linear regression analysis further demonstrated a strong time-dependent correlation between actin cable shortening and DNMT3a nuclear localization (Figure 2F). Finally, Jasplakinolide pretreatment, which stabilizes actin filaments, abrogated Blebbistatin-induced DNMT3a nuclear localization (Figure 2C and 2E), confirming that actin remodeling is both necessary and sufficient for this process.

### ERK inactivation upon actin remodeling is responsible for nuclear localization of DNMT3a

Given that actin remodeling upon wounding and Blebbistatin treatment led to consistent nuclear localization of DNMT3a, we investigated the biochemical signal that can link these two phenomena. Actin filaments serve as a platform for ERK activation,^29^ and disruption of the actin cytoskeleton negatively regulates ERK activity.^29–32^ Interestingly, studies have suggested a strong correlation between ERK activity and cytoplasmic localization of DNMT3a.^33,34^ We therefore hypothesized that inactivation of ERK would lead to the nuclear localization of DNMT3a in differentiated keratinocytes. To test the hypothesis that ERK activity correlates with actin remodeling induced by Blebbistatin, we assessed phospho-ERK1/2 levels over the same treatment timeline used in the kinetic study of actin remodelling and nuclear localization of DNMT3a (Figure 2C, 2D and 2E). We observed that ERK activity, which is quantified as the amount of the phosphorylated protein relative to the total levels of ERK (pERK/ERK), is reduced upon Blebbistatin treatment in a time-dependent manner (Figure 3A). A significant downregulation of pERK1/2 is observed from 30 minutes post-treatment (Figure 3A), that parallels the increase in nuclear DNMT3a localization (Figure 2D and 2E), suggesting an inverse correlation between ERK activity and nuclear localization of DNMT3a. To validate if the ERK inactivation by Blebbistatin is mediated by actin remodelling, we stabilized the actin cytoskeleton with Jasplakinolide and found that it prevented Blebbistatin-induced ERK inactivation (Figure S3B). Collectively, these experiments indicate that actin remodelling leads to ERK inactivation, which is directly correlated with the nuclear localization of DNMT3a. Furthermore, stabilizing actin alone by Jasplakinolide treatment increased the pERK level significantly (Figure S3A). This is consistent with studies that suggest the role of intact actin cytoskeleton as a platform for ERK activation.^29^

**Figure 3:**
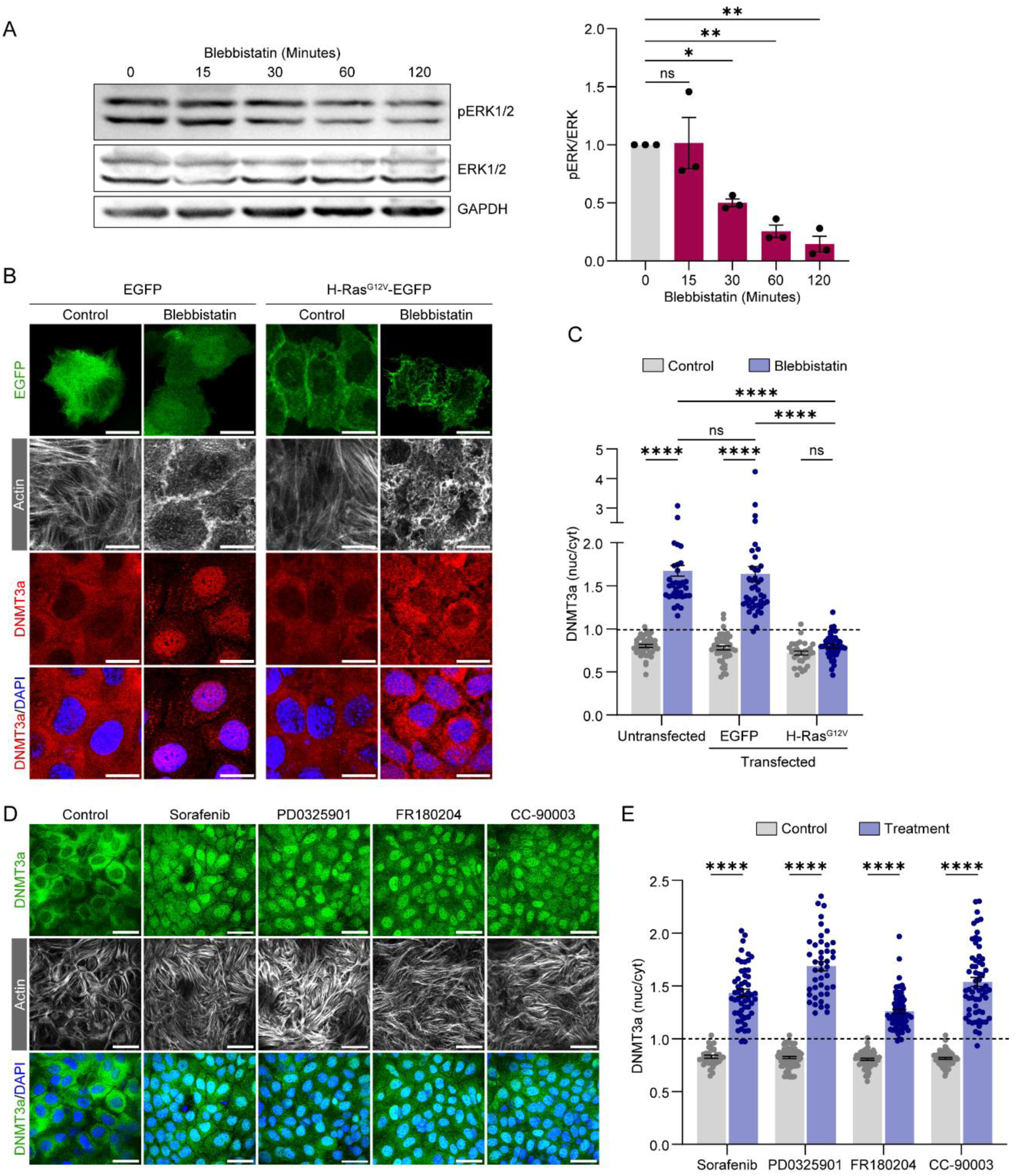
Actin remodeling-dependent ERK inactivation enables DNMT3a nuclear localization. **(A)** Western blot and quantification of pERK/ERK upon Blebbistatin treatment. **(B)** DNMT3a localization (red) with phalloidin (grey) and DAPI (blue) upon EGFP- and H-RasGV12-EGFP (green) transfected cells with or without Blebbistatin treatment. **(C)** Quantification of DNMT3a (nuc/cyt) from EGFP- and H-RasGV12-EGFP (green) transfected cells with or without Blebbistatin treatment. **(D)** DNMT3a localization (green) with phalloidin (grey) and DAPI (blue) upon treatment with ERK inhibitors. **(E)** Quantification of DNMT3a (nuc/cyt) upon treatment with ERK inhibitors. Scale bar: 20 µm for **(B)** and 50 µm for **(D)**. The data are represented as mean ± s.e.m. n=3 for **(A)**; n=4 for (B-E). P-values in **(A)**, **(C)** and **(D)** were calculated using 1-way ANOVA with Tukey’s multiple comparison test where ns= P > 0.05, *= P ≤ 0.05, **= P ≤ 0.01, ***= P ≤ 0.001, ****= P ≤ 0.0001.

To determine the necessity of ERK inactivation for the nuclear localization of DNMT3a upon actin cytoskeleton disruption, we overexpressed a constitutively active mutant of H-Ras (H-Ras^G12V^), tagged with EGFP. As expected, cells expressing EGFP alone showed cytoplasmic localization of DNMT3a under homeostasis, which dramatically shifted to the nucleus upon Blebbistatin treatment (Figure 3B and 3C). However, cells expressing H-Ras^G12V^-EGFP exhibited cytoplasmic localization of DNMT3a even upon Blebbistatin treatment (Figure 3B and 3C). Interestingly, the actin organization in the differentiated keratinocytes was comparable between EGFP and H-Ras^G12V^-EGFP overexpressing cells upon Blebbistatin treatment. Thus, the maintenance of the ERK activity in the H-Ras^G12V^-EGFP overexpressed cells led to cytoplasmic localization of DNMT3a despite the fact that the actin was disrupted (Figure 3B and 3C). This suggests that downregulation of ERK activity upon wound-induced actin cytoskeleton disruption is necessary for the nuclear localization of DNMT3a.

To understand if ERK inactivation is sufficient to stimulate the nuclear translocation of DNMT3a, we examined the effect of pharmacological inhibitor of ERK activity on the subcellular localization of DNMT3a (Figure S3C). Differentiated epidermal keratinocytes were treated with inhibitors of upstream regulators of ERK activity, namely, Sorafenib (Raf inhibitor); and PD0325901 (MEK inhibitor), leading to ERK inactivation (Figure S3D and S3E) and the subcellular localization of DNMT3a was examined. Merely treating the cells with these inhibitors was sufficient to induce nuclear localization of DNMT3a (Figure 3D and 3E). To further validate this, we have treated cells with specific ERK inhibitors - FR180204 (catalytic inhibitor) and CC-90003 (allosteric inhibitor). Inhibition of ERK activity by these specific inhibitors showed predominant nuclear localization of DNMT3a (Figure 3D and 3E). Importantly, inhibition of ERK activity led to nuclear localization of DNMT3a without any noticeable change in the actin cytoskeleton (Figure 3D), confirming that the status of ERK activity alone determines the subcellular localization of DNMT3a.

To study the relevance of ERK inactivation-induced nuclear location of DNMT3a in the wounding context, we examined the mean fluorescence intensity (MFI) of pERK1/2 upon *in vitro* scratch wounding. Upon quantification, we found that pERK MFI does not change in 0-and 2-minute post-wounding, where DNMT3a localizes in the cytoplasm (Figure S3F and S3G). However, there is a significant reduction in pERK MFI in 5-, 10-, and 15-minute post-wounding, which coincides with the wound-induced nuclear localization of DNMT3a. Total ERK level, however, does not change throughout the wounding timeline (Figure S3F, S3H), suggesting that it is the ERK activity rather than expression that dictates the subcellular localization of DNMT3a. Collectively, these data suggest that wound-induced actin remodelling results in ERK inactivation, which in turn drives the nuclear translocation of DNMT3a.

### MEK and DUSP6 orchestrate the nuclear localization of DNMT3a by inactivating ERK1/2

ERK inactivation can occur via two mechanisms: (a) downregulation of upstream MEK activity and/or (b) upregulation of ERK-specific phosphatase activity. To assess these possibilities during actin remodeling, we first examined MEK activity upon Blebbistatin treatment across the same time course that produced ERK inactivation and DNMT3a nuclear localization. We found that Blebbistatin treatment caused a reduction in MEK activity at early time points (15 and 30 min), followed by a significant increase at later time points (60 and 120 min) (Figure 4A). To determine whether the early decrease in MEK activity was attributable to Blebbistatin-induced actin disruption, cells were treated with Jasplakinolide prior to Blebbistatin using the same regimen that abrogated Blebbistatin-induced actin disruption (Figure 2C and S2E), ERK inactivation (Figure S3B), and DNMT3a nuclear localization (Figure 2C and 2E). Stabilization of actin by Jasplakinolide prevented Blebbistatin-induced MEK inactivation (Figure S4A), indicating that actin remodeling directly regulates MEK activity during this early phase. While the initial decrease in MEK activity (15 and 30 minutes) could explain the downregulation of pERK levels, increase in the MEK activity in the later time points (60 and 120 minutes) suggest that there could be another mechanism involved in the downregulation of pERK specifically in the later timepoints.

**Figure 4:**
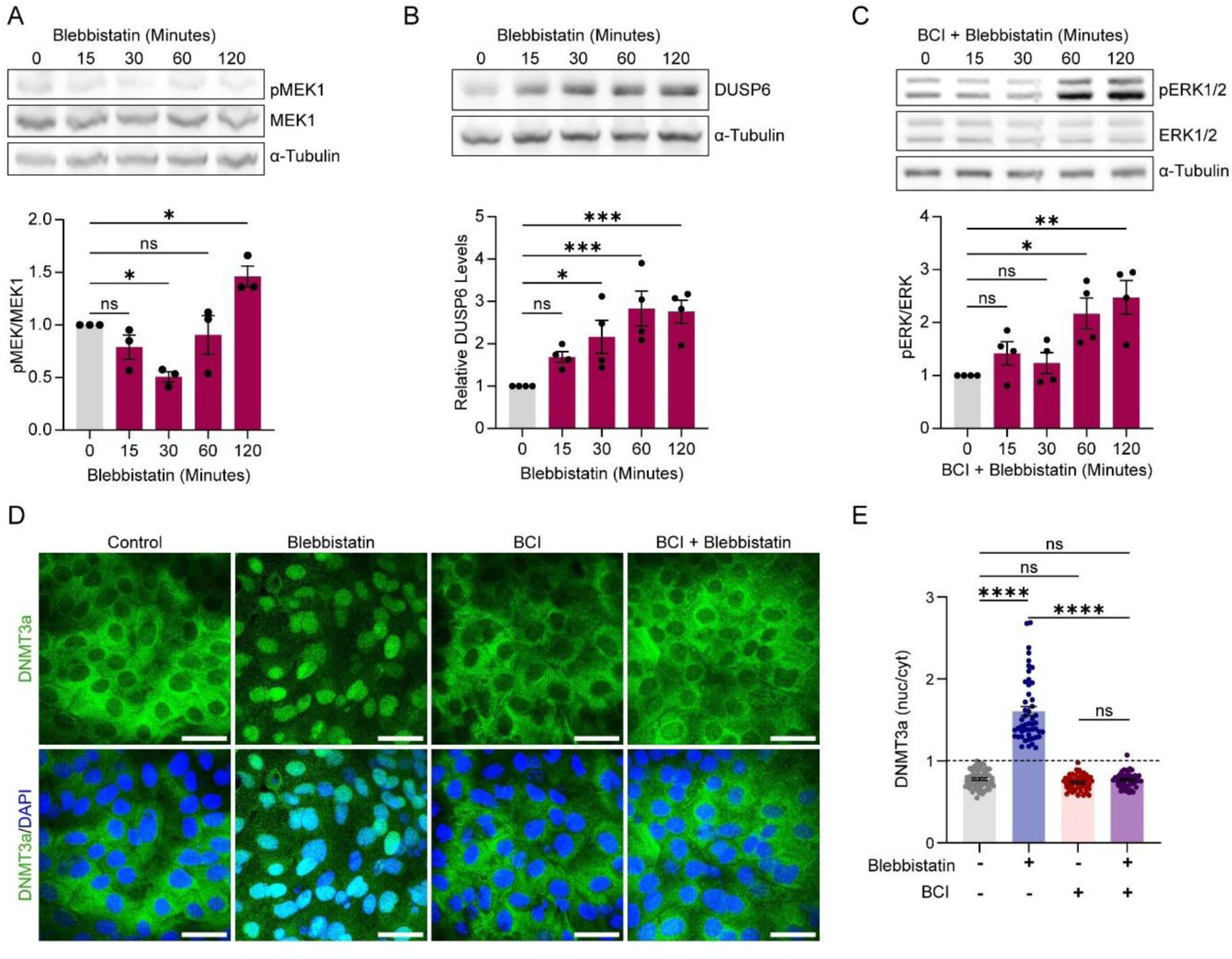
Coordinated regulation of ERK1/2 by MEK and DUSP6 drives DNMT3a nuclear localization. **(A)** Western blot and quantification of pMEK1/MEK1 upon Blebbistatin treatment. **(B)** Western blot and quantification of DUSP6 upon Blebbistatin treatment. **(C)** Western blot and quantification of pERK/ERK upon BCI + Blebbistatin treatment. **(D-E)** DNMT3a localization (green) with DAPI (blue) and quantification of DNMT3a (nuc/cyt) upon Blebbistatin, BCI, and BCI + Blebbistatin treatment. Scale bar: 50 µm for **(D)**. The data are represented as mean ± s.e.m. n=3 for **(A)**; n=4 for **(B-E)**. P-values in **(A-C)** and **(E)** were calculated using 2-way ANOVA with Tukey’s multiple comparison test, where ns= P > 0.05, *= P ≤ 0.05, **= P ≤ 0.01, ***= P ≤ 0.001, ****= P ≤ 0.0001.

This led us to hypothesize that ERK inactivation at the later timepoints could be due to the upregulation of an ERK-specific phosphatase upon actin disruption. One such ERK-specific phosphatase, well-characterized to regulate this cytoplasmic kinase in epidermal keratinocytes, is Dual Specific Phosphatase 6 (DUSP6) or MAP Kinase Phosphatase 3 (MKP3).^35,36^ We first examined if actin remodelling by Blebbistatin has any effect on DUSP6 expression. To this end, we profiled DUSP6 level upon Blebbistatin treatment, following the same treatment regimen used to profile ERK and MEK activity. Western blot data revealed an increase in expression of DUSP6 at the protein level as early as 30 minutes after Blebbistatin treatment (Figure 4B). We then investigated whether DUSP6 is involved in the downregulation of ERK activity upon actin remodelling. We treated cells with the DUSP6 inhibitor, BCI followed by Blebbistatin and observed a prevention of the early ERK inactivation (Figure 4C). Interestingly, at later timepoints (60- and 120-minutes) this treatment regimen resulted in a heightened activity of ERK (Figure 4C). The inhibition of DUSP6 also resulted in a concomitant cytoplasmic localization of DNMT3a even in the presence of Blebbistatin (Figure 4D and 4E). These results suggest an active role of DUSP6 in the regulation of ERK activity specifically in the later timepoints of Blebbistatin-induced actin remodeling. Under homeostasis, BCI treatment alone substantially increased the pERK levels as early as 15 minutes and remained elevated throughout the treatment regimen (Figure S4B). This is consistent with earlier reports documenting the cycling of ERK in epidermal keratinocytes between an “off” and “on” mode^35^. Collectively, these data suggests a bimodal regulatory process involving two key players – the upstream kinase, MEK and a cytoplasmic phosphatase, DUSP6. The early response (15 to 30 minutes) of ERK inactivation is due to the downregulation of MEK activity. Whereas the later maintenance of ERK inactivation (60 to 120 minutes) is due to the phosphatase activity of DUSP6.

In the context of wounding healing, inhibition of DUSP6 also prevented ERK inactivation (Figure S4C and S4D). This failure to inactivate ERK by BCI treatment also resulted in the cytoplasmic retention of DNMT3a even under wounding conditions (Figure S4C and S4E).

### Phosphorylation at S255 enhances intra-molecular contacts with the NLS, potentially masking its accessibility

It has been reported that DNMT3a contains a functional nuclear localization signal (NLS, 197-<u>KR</u>DEWLARWKREAE<u>KK</u>A<u>K</u>-214 of human DNMT3a) within its N-terminal domain,^37^ as indicated in the schematic (Figure 5A). Consistent with this, we observed that the N-terminal domain of hDNMT3a (N), when expressed as an EGFP fusion in keratinocytes, exhibited distinct nuclear localization even under homeostatic conditions. In contrast, the full-length hDNMT3a–EGFP (FL) localized in the cytoplasm along with nucleus (Figure 5B) but translocated to the nucleus upon Blebbistatin or MEK inhibitor treatment (Figure 5F and 5G). These observations suggest the presence of a regulatory mechanism that masks an otherwise competent NLS under homeostatic conditions.

**Figure 5:**
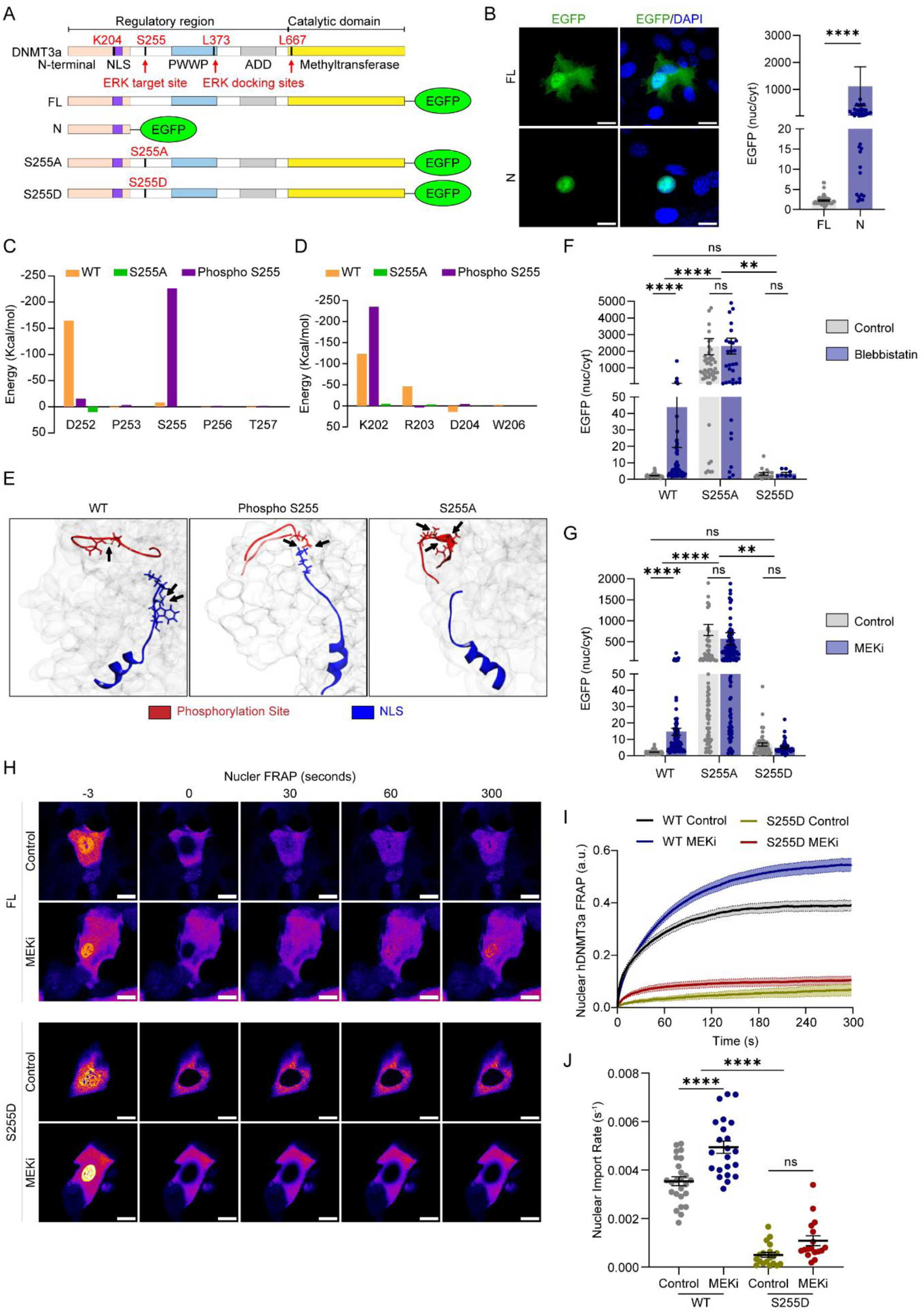
Phosphorylation of DNMT3a at S255 mediates intramolecular masking of the NLS. **(A)** Domain map of DNMT3a with NLS and critical residues for ERK docking and targeting sites. **(B)** Subcellular distribution and quantification of EGFP-tagged full length hDNMT3a (FL) and the N-terminal domain of hDNMT3a (N) containing the NLS. **(C)** and **(D)** Non-bond interaction analysis and comparison of the critical residues of DNMT3a among WT, phospho-mutant (S255A) and phospho-S255 structures at the ERK targeting site (S255) and NLS respectively. **(E)** Hydrogen-bond interaction analysis between ERK targeting site (S255) (Red) and NLS (Blue) and comparison of the bonding patterns among WT, phospho-mutant (S255A) and phospho S255 structures. Hydrogen bonds are marked in black arrows. **(F-G)** Quantification of nucleocytoplasmic localization of EGFP-tagged full length wild type hDNMT3a (WT), phospho-mutant (S255A) and phospho-mimetic (S255D) upon Blebbistatin and MEK inhibitor (PD0325901) treatment respectively. **(H)** Representative frames from the FRAP experiments. The nucleus was photobleached at t = 0s in EGFP-tagged WT and S255D expressing cells under homeostasis or Blebbistatin treatment. **(I)** Quantification of nuclear fluorescence after photobleaching the nucleus in EGFP-tagged WT and S255D expressing cells under homeostasis or MEK inhibition. **(J)** Quantification of the nuclear import rate of EGFP-tagged WT and S255D under homeostasis or MEK inhibition. Scale bar: 20 µm for all images. The data are represented as mean ± s.e.m. n=4. P-values in **(B)** calculated using unpaired Welch’s t test, in **(F-G)** and **(J)** were calculated using two-way ANOVA with Tukey’s multiple comparison test where ns= P > 0.05, *= P ≤ 0.05, **= P ≤ 0.01, ***= P ≤ 0.001, ****= P ≤ 0.0001.

To investigate whether ERK-mediated phosphorylation could regulate NLS accessibility in DNMT3a, we generated a structural model of full-length hDNMT3a using iTASSER server^38^ and examined the conformational consequences of mutating the ERK target site (S255) reported in a previous study.^39^ Structural comparison with available DNMT3a domain structures revealed high overall similarity, supporting the suitability of the model for mechanistic analysis (Figure S5A). Molecular dynamics analyses revealed that wild-type and phospho-S255 DNMT3a rapidly stabilized and reached conformational equilibrium, whereas the S255A mutant exhibited pronounced structural fluctuations, consistent with reduced conformational stability (Figure S5B). These findings suggest that phosphorylation at S255 contributes to structural integrity of the protein. Importantly, root mean square fluctuation (RMSF) analysis revealed that ERK1/2 interaction potential is likely preserved in all three protein states (Figure S5C). Instead, structural perturbations were localized to the region encompassing the phosphorylation loop (S255) and the N-terminal nuclear localization signal (NLS) (Figure S5C). Detailed interaction mapping revealed that residues surrounding S255 (D252, P253, S255, P256 and T257) engage in an extensive intramolecular interaction network with key NLS residues (K202, R203, D204 and W206) (Figure 5C and 5D; Supplementary Table 1 and 2). Phosphorylation at S255 substantially strengthened electrostatic interactions with K202 while weakening contacts with D252, resulting in a redistribution of intramolecular forces that enhanced coupling between the phosphorylation loop and the NLS (Figure 5C and 5D). This interaction pattern was absent in the S255A mutant, which lacked stable contacts between these regions (Figure 5C and 5D). Consistent with these findings, hydrogen bond analysis demonstrated that in the unphosphorylated protein, interactions were confined within the phosphorylation site and within the NLS, with no direct coupling between the two regions (Figure 5E). In contrast, phosphorylation at S255 induced stable hydrogen bonds linking pS255 directly to K202 within the NLS, effectively bridging the two regions (Figure 5E). The S255A mutant failed to establish such cross-region interactions and instead displayed altered local bonding confined to the phosphorylation loop, consistent with loss of NLS masking (Figure 5E).

To experimentally validate these *in silico* predictions, we generated phospho-mutant (S255A) and phospho-mimetic (S255D) variants of full-length hDNMT3a fused to EGFP and examined their subcellular localization in keratinocytes. Under homeostatic conditions, WT hDNMT3a–EGFP was expressed throughout the cell but completely translocated to the nucleus upon Blebbistatin treatment (Figure 5F). In contrast, the phospho-mutant S255A exhibited total nuclear localization even under homeostasis. However, the phospho-mimetic S255D mutant localized in the cytoplasm of the even in the presence of Blebbistatin (Figure 5F). Together, these results indicate that phosphorylation of DNMT3a at S255 plays a critical regulatory role in controlling its nuclear localization. These results were similar when cells were treated with a MEK inhibitor (Figure 5G).

To confirm these results, we employed fluorescence recovery after photobleaching (FRAP) to directly quantify the nuclear import dynamics of the EGFP-tagged hDNMT3a variants. Following complete photobleaching of the nuclear compartment, nuclear fluorescence recovery was monitored up to 300 seconds under homeostatic conditions and upon MEK inhibition using PD0325901. WT hDNMT3a exhibited a basal level of nuclear import following photobleaching, indicating continuous and dynamic nucleocytoplasmic shuttling to maintain steady-state equilibrium (Figure 5H, 5I and 5J). Importantly, MEK inhibition significantly enhanced the nuclear import rate of WT hDNMT3a, consistent with a shift in this equilibrium upon ERK inactivation. In contrast, the phospho-mimetic S255D variant showed markedly reduced nuclear import following photobleaching, even in the presence of PD0325901 (Figure 5H, 5I and 5J), indicating that phosphorylation at S255 is sufficient to restrain DNMT3a nuclear entry.

Together, these results indicate that phosphorylation of DNMT3a at S255 promotes intramolecular coupling between the phosphorylation site and the NLS, leading to NLS masking and cytoplasmic retention. Conversely, loss of S255 phosphorylation disrupts this coupling, resulting in NLS exposure and enabling nuclear translocation, providing a structural basis for the phosphorylation-dependent regulation of DNMT3a localization observed in cells.

## Discussion

Studies on the mechanisms underlying wound healing have traditionally focused on biochemical wound cues, including PAMPs and DAMPs.^18,40,41^ However, increasing evidence indicates that wound-induced mechanical cues also play a critical role in orchestrating the cell-state transition required for tissue repair.^17,23,42^ Previously we observed that mechanical cues can regulate the subcellular localization of DNMT3a in the wound response,^17,43,44^ but the mechanism remained unclear. We demonstrate that mechanical disruption of the actin cytoskeleton upon wounding leads to ERK inactivation, which in turn promotes the nuclear localization of DNMT3a (Figure S6). This translocation is regulated through post-translational modification at the serine 255 (S255) residue, where ERK-dependent phosphorylation appears to mask the nuclear localization signal (NLS) of DNMT3a. These findings reveal a previously unrecognized mechanotransduction pathway by which cytoskeletal integrity and ERK signaling modulate the subcellular distribution of an epigenetic regulator during tissue injury.

DNMT3a has been shown to undergo dynamic nucleo-cytoplasmic localization in development as well as disease contexts. For example, its localization shifts from the nucleus to the cytoplasm during the 2-and 4-cell stage of embryonic development.^45,46^ In glomerular mesangial cells, cytoplasmic localization is induced upon exposure to high glucose thereby alleviating diabetic nephropathy.^34^ In cardiac fibroblasts, hypoxia leads to increased expression and nuclear localization of DNMT3a.^47^ These reports suggest that this dynamic nucleo-cytoplasmic shuttling of DNMT3a can induce global transcriptional regulation enabling rapid cellular adaptation - suggesting another layer of regulation of an important epigenetic regulator. In addition to our findings, there have been complementary reports demonstrating that mechanical stimuli can alter DNMT3a localization. For instance, mechanical cues from denatured collagen matrices promoted nuclear localization of DNMT3a in smooth muscle cells.^33^

Transcription factors and epigenetic regulators typically harbor nuclear localization signals (NLSs) and nuclear export signals (NESs) that govern their subcellular distribution through coordinated recognition by the nuclear import–export machinery.^48,49^ DNMT3a contains one experimentally characterized NLS within its N-terminal domain^37^ and two additional putative NLS motifs within its regulatory region. Despite the presence of these NLS motifs, DNMT3a frequently exhibits cytoplasmic localization.^17,33,34,45,46^ This paradox suggests additional layers of regulation that restrict its nuclear accumulation. Although DNMT3a contains a putative nuclear export signal (NES) within its structurally compact catalytic domain (Figure S5A), its structural context makes dynamic regulation improbable. Classical leucine-rich NESs are typically constitutive and only rarely subjected to rapid regulatory control, in contrast to nuclear localization signals that often reside in flexible regions amenable to post-translational masking.^48,50^ Such masking mechanisms, often mediated by phosphorylation, enable rapid and reversible control of nuclear import without altering protein abundance.^50^ In the case of DNMT3a, our findings reveal a negative regulatory mechanism whereby phosphorylation of a neighboring serine residue (S255) masks the NLS, preventing nuclear entry under homeostatic conditions.

The dynamic subcellular partitioning of DNMT3a is reflected in the cyclical activity of ERK in epidermal keratinocytes, thus hinting at the connection between these two phenomena. Temporal mapping of MAPK using a genetic activity reporter demonstrated pulsatile ERK activity in the epidermis.^35^ These oscillatory dynamics are maintained by a feedback network involving the dual-specificity phosphatases DUSP6 and DUSP10, which fine-tune ERK activity to preserve epidermal homeostasis.^35^

We propose that under homeostatic conditions, intact actin cytoskeleton sustains ERK activity through continuous input from the Ras-Raf-MEK cascade. Upon wounding, cytoskeletal disruption relieves intracellular tension, leading to activation of DUSP6 and consequent ERK dephosphorylation. This transition toward ERK inactivity functions as a signaling switch that unmasks the nuclear localization signal of DNMT3a, enabling its nuclear translocation and initiation of epigenetic reprogramming. Together, these findings provide a mechanistic framework linking mechanical disruption to kinase-phosphatase balance and chromatin regulation, explaining how local loss of cytoskeletal tension can rapidly remodel the epigenetic landscape during tissue repair.

Finally, our work raises important implications for pathological wound healing. Diabetic skin has been reported to exhibit altered mechanical properties, including weakened biomechanical strength and cytoskeletal disorganization, which may affect wound responsiveness.^51,52^ These changes may lead to chronic misregulation of ERK-DNMT3a signaling, either through inappropriate basal ERK suppression or failure to dynamically modulate DNMT3a localization during injury. Whether DNMT3a remains aberrantly nuclear or cytoplasmic in diabetic wounds, and how this impacts epigenetic plasticity and repair competence, represents an important avenue for future study.

## Materials and Methods

### Reagents and Tools Table

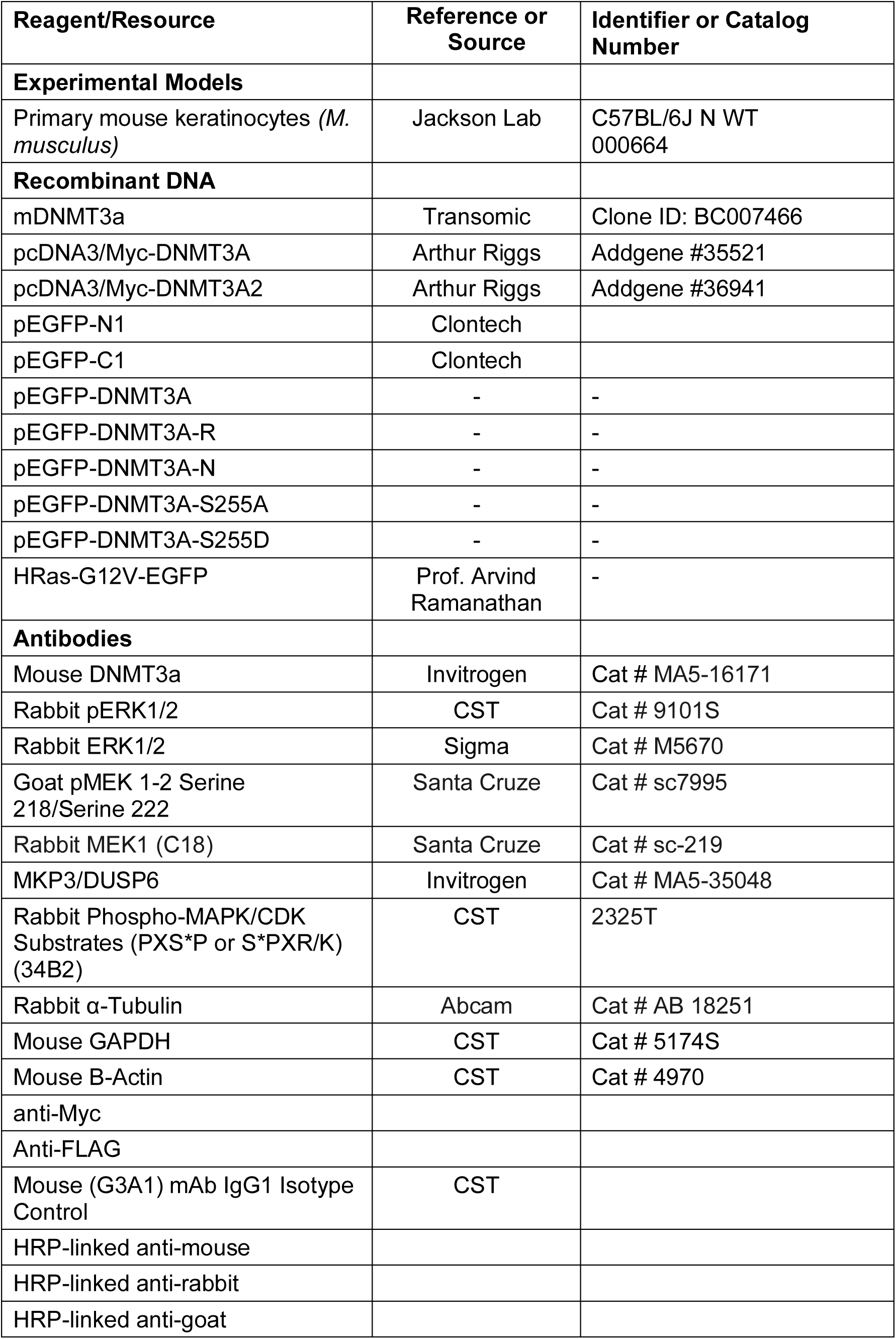
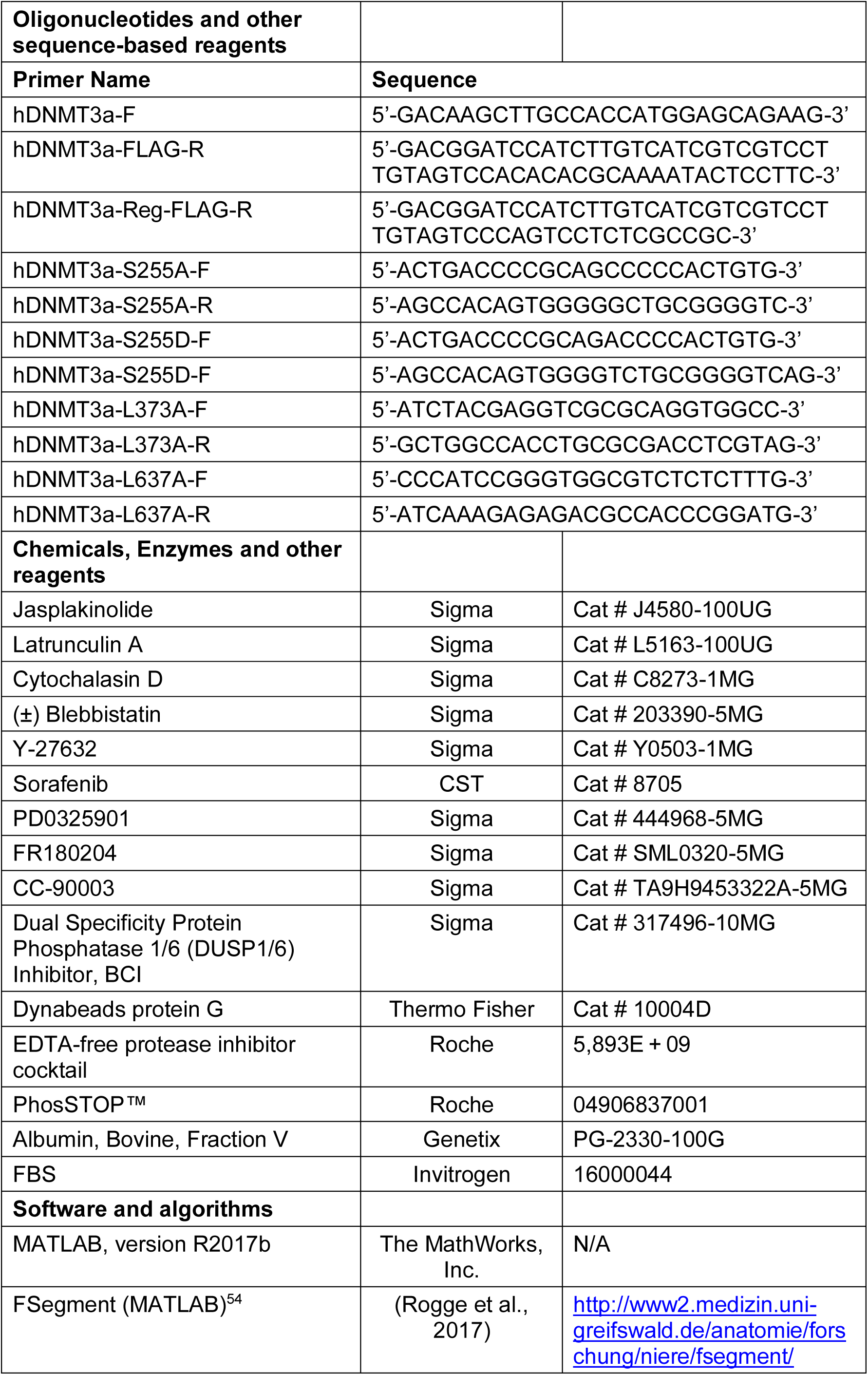
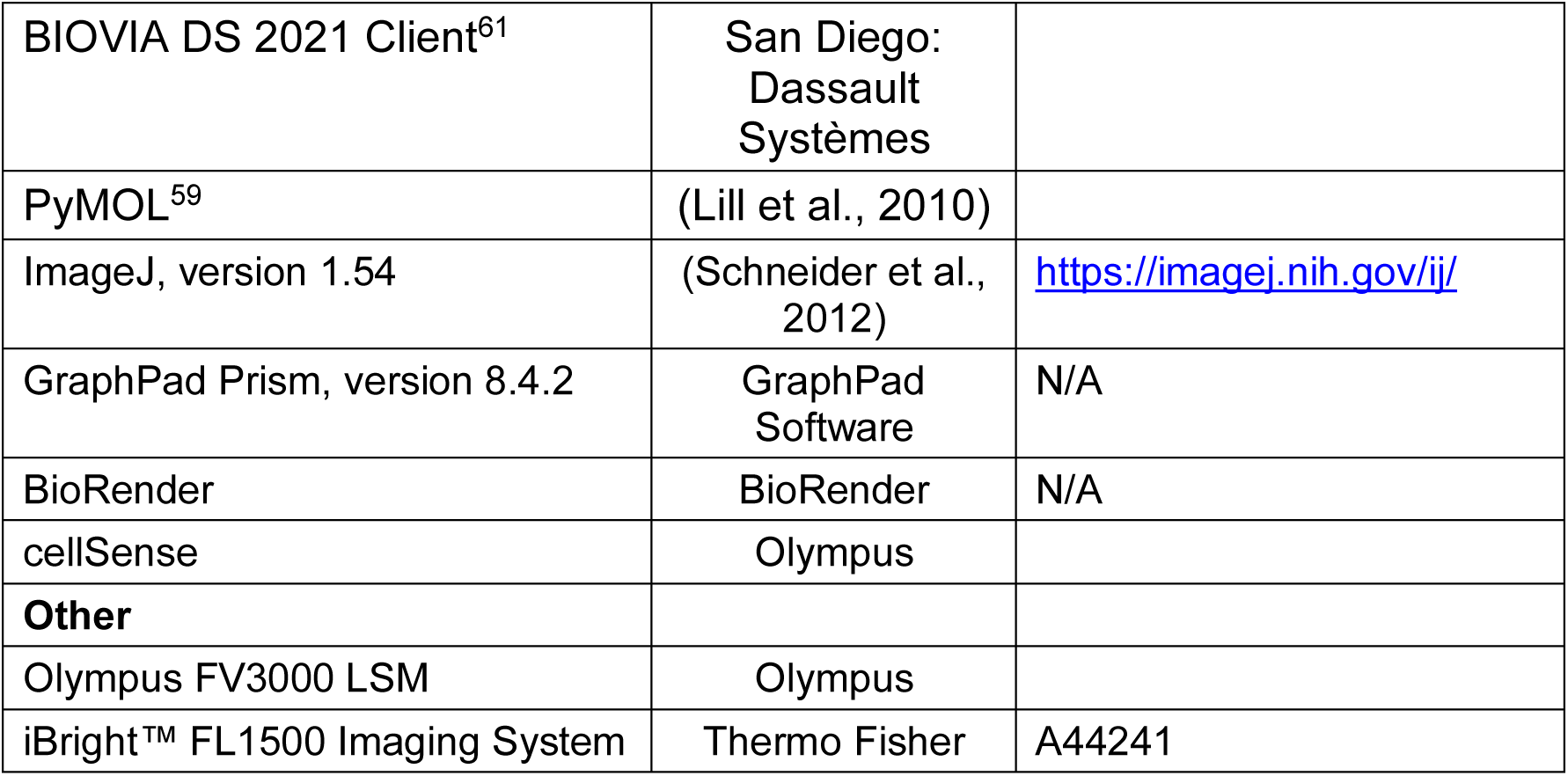

### Primary keratinocyte culture and differentiation

Primary neonatal mouse epidermal keratinocytes were isolated from P0-P3 pups and cultured as described elaborately in the protocol by Nowak and Fuchs ^53^. Briefly, the piece of skin tissue was incubated in dispase at 37°C for 1h. With the help of fine forceps, the epidermis layer was separated from the dermis layer and placed inside a 3.5 cm culture dish containing 1 ml of trypsin: versene (1:1 ratio) for 10 mins at room temperature. Following that, loosen-up cells were taken out in 4 ml of low Ca^2+^ (0.05 mM) E-media. Further, they were filtered through a 70 mm membrane, spun down, and re-suspended in fresh low Ca^2+^ E-media. Finally, they were plated on pre-cultured feeder cells (3T3-J2 fibroblasts at 70-80% confluence) in low Ca^2+^ E-media for subsequent culture till 10-12 passages, when they became feeder-free. For differentiation of these primary keratinocytes, high Ca^2+^ (1.2 mM) E-media was added to the culture dish and was incubated for 72 hrs.

### Plasmids and transfection

Plasmid transfections in primary mouse keratinocytes were performed with Lipofectamine™ 3000 Reagent (ThermoFisher, #L3000015) following manufacturer’s protocol.

### Western blot analysis

The cell lysates were prepared in RIPA buffer (50 mM Tris-Cl, pH 7.5, 150 mM NaCl, 1% NP40) with protease inhibitor cocktail (50X PIC) (Sigma, #P2714) and the sample was sonicated at 4°C. Sonicated protein lysate was then mixed with 5X Laemmle sample loading buffer and heated at 96°C for 10 mins before running on the polyacrylamide gel followed by nitrocellulose membrane transfer. After the transfer was done, to get rid of noise, the membrane was blocked using either 5% Bovine Serum Albumin (BSA) in Tris-Cl buffer Saline containing 0.1% Tween (TBST) or 5% Blotto, fat-free milk (Santa Cruz Biotechnology, sc-2325) upon solubilizing in same TBST buffer for one hour. After blocking, the blots were kept for either 2 hours at room temperature or overnight at 4°C incubation with primary antibodies of recommended concentration in the blocking buffer. After the primary antibody incubation was over, the blots were washed thoroughly with TBST buffer followed by HRP-conjugated secondary antibody incubation for half an hour at room temperature followed by vigorous washing in TBST. Signals were detected using western blot developing solution, Enhanced Chemiluminescence substrate (ECL, Merck) using the iBright FL (Thermo) detector.

### Immunofluorescence staining

Cells were cultured on glass coverslips coated with collagen or fibronectin and differentiated. After the treatment, cells were fixed using 4% PFA for 5-10 min followed by washing with 1X TBS. After washing cells were permeabilized with 0.2% Triton X100 (in 1X TBS) for 15 minutes followed by washing with 1X TBS and blocking for one hour with IF blocking solution. After blocking, they were incubated with primary antibody for 2 hours at room temperature or overnight at 4°C. The next day, the coverslips were again washed and probed with fluorophore-conjugated secondary antibodies at room temperature for 30 minutes, and images were taken using Olympus FV3000 confocal microscope using PLAPON 60X oil objective. Acquired images were further analyzed using FIJI (ImageJ software). The primary antibodies and their respective dilutions used for immunofluorescence staining are as follows: DNMT3a (Invitrogen, Cat # MA5-16171; 1:200), Phospho ERK1/2 (CST, Cat # 9101S; 1:1000), Total ERK (Sigma, Cat # M5670; 1:10000), a-Tubulin (Abcam, Cat # AB 18251; 1:4000), GAPDH (CST, Cat # 5174S; 1:4000), b-Actin (CST, Cat # 4970; 1:4000), pMEK 1-2 Serine 218/Serine 222 (Santa Cruze, Cat # sc7995; 1:1000), Rabbit MEK1 (C18) (Santa Cruze, Cat # sc-219; 1:1000), MKP3/ DUSP6 (Invitrogen, Cat # MA5-35048; 1:1000). Alexa Fluor™ 488, 561 and 647-labelled secondary antibodies (Invitrogen) were used at a dilution of 1:400. Alexa Fluor™ 488. 568 and 647 Phalloidin (Invitrogen, Cat # A12379, A12380 and A22287) were used to probe for the actin cytoskeleton. Hoechst (Invitrogen, Cat # H1399) or DAPI (Sigma, Cat # MBD0015-5ML) were used to mark the nuclei.

### DNMT3a (nuc/cyt) Analysis

The ratio of the mean immunofluorescence intensity of DNMT3A from the nucleus/cytoplasm was calculated following the strategy developed by *Nardone* et al. using the following formula.

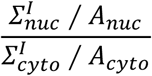

where 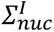 and 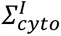 represent the sum of the raw integrated density for the pixels in the nuclear and cytoplasmic region respectively, and 𝐴_𝑛𝑢𝑐_ and 𝐴_𝑐𝑦𝑡𝑜_ represent the area of the corresponding regions.

Since differentiated keratinocytes form a layer of cells tightly adhered together by cellular junctions, we have considered individual confocal image fields for the calculation of average DNMT3a (nuc/cyt), 𝑅_𝑁𝐶_ using the strategy followed by Zhang et al., 2021 using the following modified formula^47^:

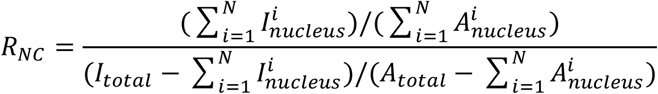

where 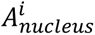 is the area of the nucleus *i* in the zone containing *N* nuclei, 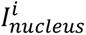 is the total fluorescence of nucleus *i*, 𝐼_𝑡𝑜𝑡𝑎𝑙_ is the total fluorescence intensity of the confocal image field, and 𝐴_𝑡𝑜𝑡𝑎𝑙_is the total area of the confocal image field. Each data point on every bar graph represents the mean DNMT3a (nuc/cyt) from individual confocal frames containing approximately 50-70 cells.

### Actin Quantification

Actin cytoskeleton was visualized by fluorophore tagged phalloidin (Alexa Fluor™ 488, 568 or 647) probes using Olympus FV3000 confocal microscope. The actin cable length under different conditions were then analyzed using MATLAB based tool FSegment developed by Rogge et al.^54^ Briefly, actin cables were segmented using a trace algorithm applying the parameters for detection of actin cables on confocal images with subtracted background noise. Segmented linear actin cables were then analyzed to quantify for individual and mean actin cable length, actin cable width and total numbers, etc. The individual actin cable length values from the confocal images across different technical and biological replicates were then represented as scatter plots or frequency distribution heatmaps.

### Fluorescence recovery after photobleaching (FRAP)

FRAP was performed to quantify nuclear import dynamics of hDNMT3a-EGFP under different genetic (WT, S255D) and pharmacological conditions. Cells expressing hDNMT3a-EGFP were imaged on an inverted confocal system using a 60× oil immersion objective. A circular region covering the entire nucleus was selected and photobleached using a 488-nm laser pulse. Time-lapse imaging was initiated immediately after bleaching, and images were acquired at high temporal resolution for the initial recovery phase. The bleach frame was defined as t = 0 s. FRAP analysis was performed using FRAP module of the Cell Sense software (Olympus).

### 3D Modeling and Structural Evaluation of DNMT3a

The human DNMT3a sequence in FASTA format was retrieved from the UniProtKB database and used as the target for structural modeling via the I-TASSER server. I-TASSER (Iterative Threading ASSEmbly Refinement) is a hierarchical method for protein structure prediction and functional annotation that integrates structural information from known protein templates.^38^ The modeling pipeline begins with the identification of appropriate structural templates from the Protein Data Bank (PDB) using the LOMETS meta-threading algorithm, followed by iterative assembly simulations of template-based fragments to generate full-length atomic models.^38^ The resulting models were evaluated for structural quality and stability using multiple validation tools. Stereochemical parameters were assessed using the Ramachandran plot implemented in PROCHECK,^55^ while non-bonded atomic interactions were examined using ERRAT.^56^ The compatibility of the modeled structure with its amino acid sequence was evaluated with Verify3D,^57^ and overall model quality was analyzed using ProSA.^58^ Refined structural models were obtained using the GalaxyRefiner platform. Root mean square deviation (RMSD) values for model superimposition were calculated in PyMOL.^59^ Post-translational modification was incorporated into the structure using the PyMOL extension,^60^ and the S255A mutant of DNMT3a was generated using the BIOVIA mutation builder with default parameters.^61^

### Molecular Dynamics (MD) Simulation

MD simulations were conducted using BIOVIA DS 2021 Client to investigate intramolecular interactions over the simulation timescale.^61^ The CHARMm force field was applied to the protein structure prior to solvation in an explicit periodic boundary model. An orthorhombic solvation cell was used, maintaining a minimum distance of 30 Å between the solute and the periodic boundary. An initial 10 ps standard dynamics cascade was performed to prepare the system for production runs. This included steepest descent minimization (2000 steps) followed by conjugate gradient minimization (2000 steps). The system was then gradually heated from 50 K to 300 K over 4 ps, equilibrated for 10 ps at 300 K, and subjected to a 10 ps pre-production run under NPT conditions with automatic electrostatics. Subsequently, production MD simulations were executed on a GPU for 100,000 ps (100 ns) using the restart file generated from the cascade step. MD trajectories were recorded every 0.5 ns, and energy profiles were extracted for further analysis.

### QM/MM Calculations

QM/MM calculations were carried out to evaluate the energetic contributions of specific residues using a hybrid quantum mechanics/molecular mechanics (QM/MM) approach in BIOVIA DS 2021 Client.^61^ Selected region is treated as quantum mechanically (CHARMm) and the remainder of the system is handled with molecular mechanics (DMol³) A non-bonded interaction pair distance cutoff of 14 Å was applied, and DMol³ settings were kept at default parameters.

The QM/MM interaction energy was calculated as the sum of two components:

1. **QM/MM Electrostatic Interaction Energy** – the Coulombic interaction energy between the electron density of the QM atoms and the point charges of the MM atoms. This represents an approximation of the full QM/MM electrostatic interaction energy, as the energy required to polarize the QM density from its vacuum state is not included.
2. **QM/MM van der Waals Interaction Energy** – computed using the CHARMm simulation engine by applying van der Waals force field parameters to the complete system.

The total QM/MM interaction energy was obtained by summing the electrostatic and van der Waals contributions.^61^

### Statistical analysis

The data are presented as the means ± standard error of mean (SEM) from three or more biological replicates. All statistical analyses were performed in GraphPad Prism (v8.4.2). For comparison between two groups, a two-tailed unpaired Student’s t test with Welch’s correction was used. For comparisons between multiple groups, ordinary one-way ANOVA followed by Tukey’s post hoc test (parametric data with one variable) or ordinary two-way ANOVA with Sidak’s test were performed (parametric data with two independent variables).

## Author contributions

Conceptualization: JA, CJ; Methodology: JA, BD, EG, TS, AD, SK, AG, DP, CJ; Investigation: JA, BD, EG, TS, AD, SK; Formal Analysis: JA, BD, EG, AD; Visualization: JA, BD, EG, AD; Funding Acquisition: CJ, DP; Supervision: CJ; Writing - Original Draft Preparation: JA, CJ; Writing - Review and Editing: JA, EG, AG, CJ.

## Acknowledgments

The authors would like to thank members of the Jamora laboratory for critical review of the manuscript. We thank Srinivas Animireddy and Arjun Guha from BRIC-inStem for insightful discussions. This work was supported by core funds from BRIC-inStem and Shiv Nadar Institution of Eminence and by grants from the Department of Biotechnology of the Government of India (BT/PR8738/AGR/36/770/2013 and BT/PR32539/BRB/10/1814/2019) and the Indian Council of Medical Research (IIRPIG-2024-01-01387). Animals are maintained at the BLiSC Animal Care and Resource Center (ACRC) and is partially supported by the National Mouse Research Resource grant BT/PR5981/MED/31/181/2012;2013-2016;2018 and 102/IFD/SAN/5003/2017-2018 from the Department of Biotechnology. EG was supported by the funding of DST ISPIRE fellowship. We thank Central Imaging and Flow Cytometry Facility (CIFF) at the Bangalore Life Science Cluster for support with imaging and analysis.

## References

1. Lewis, C.J., Mardaryev, A.N., Sharov, A.A., Fessing, M.Y., and Botchkarev, V.A. (2014). The Epigenetic Regulation of Wound Healing. https://home.liebertpub.com/wound 3, 468–475. 10.1089/WOUND.2014.0522.

2. Lewis, C.J., Stevenson, A., Fear, M.W., and Wood, F.M. (2020). A review of epigenetic regulation in wound healing: Implications for the future of wound care. Wound Repair and Regeneration 28, 710–718. 10.1111/WRR.12838.

3. Yu, H., Wang, Y., Wang, D., Yi, Y., Liu, Z., Wu, M., Wu, Y., and Zhang, Q. (2022). Landscape of the epigenetic regulation in wound healing. Front. Physiol. 13, 949498. 10.3389/FPHYS.2022.949498.

4. Luo, G., Jing, X., Yang, S., Peng, D., Dong, J., Li, L., Reinach, P.S., and Yan, D. (2019). DNA Methylation Regulates Corneal Epithelial Wound Healing by Targeting miR-200a and CDKN2B. Invest. Ophthalmol. Vis. Sci. 60, 650–660. 10.1167/IOVS.18-25443.

5. Yan, J., Tie, G., Wang, S., Tutto, A., Demarco, N., Khair, L., Fazzio, T.G., and Messina, L.M. (2018). Diabetes impairs wound healing by Dnmt1-dependent dysregulation of hematopoietic stem cells differentiation towards macrophages. Nature Communications 2017 9:1 9, 33-. 10.1038/s41467-017-02425-z.

6. Cabanel, M., Da Costa, T.P., El-Cheikh, M.C., and Carneiro, K. (2019). The epigenome as a putative target for skin repair: the HDAC inhibitor Trichostatin A modulates myeloid progenitor plasticity and behavior and improves wound healing. Journal of Translational Medicine 2019 17:1 17, 247-. 10.1186/S12967-019-1998-9.

7. Davis, F.M., Dendekker, A., Joshi, A., Wolf, S., Moore, B., Lukacs, N., and Gallagher, K. (2020). Epigenetic Regulation of Toll-like Receptor 4 Signaling Modulates Macrophage Phenotype and Impairs Diabetic Wound Healing. J. Vasc. Surg. 72, e260. 10.1016/j.jvs.2020.04.431.

8. Karnam, K., Sedmaki, K., Sharma, P., Routholla, G., Goli, S., Ghosh, B., Venuganti, V.V.K., and Kulkarni, O.P. (2020). HDAC6 inhibitor accelerates wound healing by inhibiting tubulin mediated IL-1β secretion in diabetic mice. Biochimica et Biophysica Acta (BBA) - Molecular Basis of Disease 1866, 165903. 10.1016/J.BBADIS.2020.165903.

9. Kimball, A.S., Joshi, A., Carson, W.F., Boniakowski, A.E., Schaller, M., Allen, R., Bermick, J., Davis, F.M., Henke, P.K., Burant, C.F., et al. (2017). The Histone Methyltransferase MLL1 Directs Macrophage-Mediated Inflammation in Wound Healing and Is Altered in a Murine Model of Obesity and Type 2 Diabetes. Diabetes 66, 2459–2471. 10.2337/DB17-0194.

10. Li, X., Liu, C., Zhu, Y., Rao, H., Liu, M., Gui, L., Feng, W., Tang, H., Xu, J., Gao, W.Q., et al. (2021). SETD2 epidermal deficiency promotes cutaneous wound healing via activation of AKT/mTOR Signalling. Cell Prolif. 54, e13045. 10.1111/CPR.13045.

11. Na, J., Shin, J.Y., Jeong, H., Lee, J.Y., Kim, B.J., Kim, W.S., Yune, T.Y., and Ju, B.G. (2017). JMJD3 and NF-κB-dependent activation of Notch1 gene is required for keratinocyte migration during skin wound healing. Scientific Reports 2017 7:1 7, 6494-. 10.1038/s41598-017-06750-7.

12. Jiang, Y., Xu, X., Xiao, L., Wang, L., and Qiang, S. (2022). The Role of microRNA in the Inflammatory Response of Wound Healing. Front. Immunol. 13, 852419. 10.3389/FIMMU.2022.852419/FULL.

13. Soliman, A.M., Das, S., Abd Ghafar, N., and Teoh, S.L. (2018). Role of MicroRNA in proliferation phase of wound healing. Front. Genet. 9, 335936. 10.3389/FGENE.2018.00038/FULL.

14. Vasioukhin, V., Bauer, C., Yin, M., and Fuchs, E. (2000). Directed Actin Polymerization Is the Driving Force for Epithelial Cell–Cell Adhesion. Cell 100, 209–219. 10.1016/S0092-8674(00)81559-7.

15. Vasioukhin, V., Bowers, E., Bauer, C., Degenstein, L., and Fuchs, E. (2001). Desmoplakin is essential in epidermal sheet formation. Nat. Cell Biol. 3, 1076–1085. 10.1038/NCB1201-1076.

16. Naik, S., Larsen, S.B., Gomez, N.C., Alaverdyan, K., Sendoel, A., Yuan, S., Polak, L., Kulukian, A., Chai, S., and Fuchs, E. (2017). Inflammatory memory sensitizes skin epithelial stem cells to tissue damage. Nature 2017 550:7677 550, 475–480. 10.1038/nature24271.

17. Bhatt, T., Dey, R., Hegde, A., Ketkar, A.A., Pulianmackal, A.J., Deb, A.P., Rampalli, S., and Jamora, C. (2022). Initiation of wound healing is regulated by the convergence of mechanical and epigenetic cues. PLoS Biol. 20, e3001777. 10.1371/JOURNAL.PBIO.3001777.

18. Peña, O.A., and Martin, P. (2024). Cellular and molecular mechanisms of skin wound healing. Nature Reviews Molecular Cell Biology 2024 25:8 25, 599–616. 10.1038/s41580-024-00715-1.

19. Gopalakrishnan, S., Van Emburgh, B.O., Shan, J., Su, Z., Fields, C.R., Vieweg, J., Hamazaki, T., Schwartz, P.H., Terada, N., and Robertson, K.D. (2009). A novel DNMT3B splice variant expressed in tumor and pluripotent cells modulates genomic DNA methylation patterns and displays altered DNA binding. Molecular Cancer Research 7, 1622–1634. 10.1158/1541-7786.MCR-09-0018/357182/P/A-NOVEL-DNMT3B-SPLICE-VARIANT-EXPRESSED-IN-TUMOR.

20. Vandiver, A.R., Irizarry, R.A., Hansen, K.D., Garza, L.A., Runarsson, A., Li, X., Chien, A.L., Wang, T.S., Leung, S.G., Kang, S., et al. (2015). Age and sun exposure-related widespread genomic blocks of hypomethylation in nonmalignant skin. Genome Biol. 16, 1–15. 10.1186/S13059-015-0644-Y/FIGURES/4.

21. Nardone, G., Oliver-De La Cruz, J., Vrbsky, J., Martini, C., Pribyl, J., Skládal, P., Pešl, M., Caluori, G., Pagliari, S., Martino, F., et al. (2017). YAP regulates cell mechanics by controlling focal adhesion assembly. Nature Communications 2017 8:1 8, 1–13. 10.1038/ncomms15321.

22. Karsch, S., Büchau, F., Magin, T.M., and Janshoff, A. (2020). An intact keratin network is crucial for mechanical integrity and barrier function in keratinocyte cell sheets. Cellular and Molecular Life Sciences 77, 4397–4411. 10.1007/S00018-019-03424-7/FIGURES/7.

23. Nunan, R., Campbell, J., Mori, R., Pitulescu, M.E., Jiang, W.G., Harding, K.G., Adams, R.H., Nobes, C.D., and Martin, P. (2015). Ephrin-Bs Drive Junctional Downregulation and Actin Stress Fiber Disassembly to Enable Wound Re-epithelialization. Cell Rep. 13, 1380–1395. 10.1016/j.celrep.2015.09.085.

24. Togo, T., Krasieva, T.B., and Steinhardt, R.A. (2000). A decrease in membrane tension precedes successful cell-membrane repair. Mol. Biol. Cell 11, 4339–4346. 10.1091/MBC.11.12.4339.

25. Trepat, X., Wasserman, M.R., Angelini, T.E., Millet, E., Weitz, D.A., Butler, J.P., and Fredberg, J.J. (2009). Physical forces during collective cell migration. Nature Physics 2009 5:6 5, 426–430. 10.1038/nphys1269.

26. Bubb, M.R., Spector, I., Beyer, B.B., and Fosen, K.M. (2000). Effects of Jasplakinolide on the Kinetics of Actin Polymerization: AN EXPLANATION FOR CERTAIN IN VIVO OBSERVATIONS. Journal of Biological Chemistry 275, 5163–5170. 10.1074/JBC.275.7.5163.

27. Coué, M., Brenner, S.L., Spector, I., and Korn, E.D. (1987). Inhibition of actin polymerization by latrunculin A. FEBS Lett. 213, 316–318. 10.1016/0014-5793(87)81513-2.

28. Casella, J.F., Flanagan, M.D., and Lin, S. (1981). Cytochalasin D inhibits actin polymerization and induces depolymerization of actin filaments formed during platelet shape change. Nature 293, 302–305. 10.1038/293302A0;KWRD=SCIENCE.

29. Hirata, H., Gupta, M., Vedula, S.R.K., Lim, C.T., Ladoux, B., and Sokabe, M. (2015). Actomyosin bundles serve as a tension sensor and a platform for ERK activation. EMBO Rep. 16, 250–257. 10.15252/EMBR.201439140/SUPPL_FILE/EMBR201439140.REVIEWER_COMMENTS.PDF.

30. Kawamura, S., Miyamoto, S., and Brown, J.H. (2003). Initiation and Transduction of Stretch-induced RhoA and Rac1 Activation through Caveolae. Journal of Biological Chemistry 278, 31111–31117. 10.1074/jbc.m300725200.

31. Lim, Y. Bin, Kang, S.S., Park, T.K., Lee, Y.S., Chun, J.S., and Sonn, J.K. (2000). Disruption of actin cytoskeleton induces chondrogenesis of mesenchymal cells by activating protein kinase C-α signaling. Biochem. Biophys. Res. Commun. 273, 609–613. 10.1006/BBRC.2000.2987.

32. Müller, P., Langenbach, A., Kaminski, A., and Rychly, J. (2013). Modulating the Actin Cytoskeleton Affects Mechanically Induced Signal Transduction and Differentiation in Mesenchymal Stem Cells. PLoS One 8, e71283. 10.1371/JOURNAL.PONE.0071283.

33. Jiang, J.X., Aitken, K.J., Sotiropolous, C., Kirwan, T., Panchal, T., Zhang, N., Pu, S., Wodak, S., Tolg, C., and Bägli, D.J. (2013). Phenotypic Switching Induced by Damaged Matrix Is Associated with DNA Methyltransferase 3A (DNMT3A) Activity and Nuclear Localization in Smooth Muscle Cells (SMC). PLoS One 8, e69089. 10.1371/JOURNAL.PONE.0069089.

34. Zhang, H., Li, A., Zhang, W., Huang, Z., Wang, J., and Yi, B. (2016). High glucose-induced cytoplasmic translocation of Dnmt3a contributes to CTGF hypo-methylation in mesangial cells. Biosci. Rep. 36, 362. 10.1042/BSR20160141/56452.

35. Hiratsuka, T., Bordeu, I., Pruessner, G., and Watt, F.M. (2020). Regulation of ERK basal and pulsatile activity control proliferation and exit from the stem cell compartment in mammalian epidermis. Proc. Natl. Acad. Sci. U. S. A. 117, 17796–17807. 10.1073/PNAS.2006965117/SUPPL_FILE/PNAS.2006965117.SM03.AVI.

36. Mishra, A., Oulès, B., Pisco, A.O., Ly, T., Liakath-Ali, K., Walko, G., Viswanathan, P., Tihy, M., Nijjher, J., Dunn, S.J., et al. (2017). A protein phosphatase network controls the temporal and spatial dynamics of differentiation commitment in human epidermis. Elife 6. 10.7554/ELIFE.27356.

37. Zeng, Y., Ren, R., Kaur, G., Hardikar, S., Ying, Z., Babcock, L., Gupta, E., Zhang, X., Chen, T., and Cheng, X. (2020). The inactive Dnmt3b3 isoform preferentially enhances Dnmt3b-mediated DNA methylation. Genes Dev. 34, 1546–1558. 10.1101/GAD.341925.120/-/DC1.

38. Yang, J., and Zhang, Y. (2015). I-TASSER server: new development for protein structure and function predictions. Nucleic Acids Res. 43, W174–W181. 10.1093/NAR/GKV342.

39. Kumar, D., and Lassar, A.B. (2014). Fibroblast Growth Factor Maintains Chondrogenic Potential of Limb Bud Mesenchymal Cells by Modulating DNMT3A Recruitment. Cell Rep. 8, 1419–1431. 10.1016/J.CELREP.2014.07.038.

40. Zindel, J., and Kubes, P. (2020). DAMPs, PAMPs, and LAMPs in Immunity and Sterile Inflammation. Annual Review of Pathology: Mechanisms of Disease 15, 493–518. 10.1146/ANNUREV-PATHMECHDIS-012419-032847/CITE/REFWORKS.

41. Rai, V., Mathews, G., and Agrawal, D.K. (2022). Translational and Clinical Significance of DAMPs, PAMPs, and PRRs in Trauma-induced Inflammation. Arch. Clin. Biomed. Res. 6, 673. 10.26502/ACBR.50170279.

42. Martinelli, R., Kamei, M., Sage, P.T., Massol, R., Varghese, L., Sciuto, T., Toporsian, M., Dvorak, A.M., Kirchhausen, T., Springer, T.A., et al. (2013). Release of cellular tension signals self-restorative ventral lamellipodia to heal barrier micro-wounds. Journal of Cell Biology 201, 449–465. 10.1083/jcb.201209077.

43. Yang, Z.G., Awan, F.M., Du, W.W., Zeng, Y., Lyu, J., Wu, D., Gupta, S., Yang, W., and Yang, B.B. (2017). The Circular RNA Interacts with STAT3, Increasing Its Nuclear Translocation and Wound Repair by Modulating Dnmt3a and miR-17 Function. Molecular Therapy 25, 2062–2074. 10.1016/j.ymthe.2017.05.022.

44. Yang, B., Alimperti, S., Gonzalez, M. V., Dentchev, T., Kim, M., Suh, J., Titchenell, P.M., Ko, K.I., Seykora, J., Benakanakere, M., et al. (2024). Reepithelialization of Diabetic Skin and Mucosal Wounds Is Rescued by Treatment With Epigenetic Inhibitors. Diabetes 73, 120–134. 10.2337/DB23-0258.

45. Chen, T., Ueda, Y., Dodge, J.E., Wang, Z., and Li, E. (2003). Establishment and Maintenance of Genomic Methylation Patterns in Mouse Embryonic Stem Cells by Dnmt3a and Dnmt3b. Mol. Cell. Biol. 23, 5594–5605. 10.1128/mcb.23.16.5594-5605.2003.

46. Uysal, F., Ozturk, S., and Akkoyunlu, G. (2017). DNMT1, DNMT3A and DNMT3B proteins are differently expressed in mouse oocytes and early embryos. J. Mol. Histol. 48, 417–426. 10.1007/s10735-017-9739-y.

47. Zhang, C., Zhu, H., Ren, X., Gao, B., Cheng, B., Liu, S., Sha, B., Li, Z., Zhang, Z., Lv, Y., et al. (2021). Mechanics-driven nuclear localization of YAP can be reversed by N-cadherin ligation in mesenchymal stem cells. Nature Communications 2021 12:1 12, 1–13. 10.1038/s41467-021-26454-x.

48. Lange, A., Mills, R.E., Lange, C.J., Stewart, M., Devine, S.E., and Corbett, A.H. (2007). Classical nuclear localization signals: Definition, function, and interaction with importin α. Journal of Biological Chemistry 282, 5101–5105. 10.1074/jbc.R600026200.

49. Marfori, M., Mynott, A., Ellis, J.J., Mehdi, A.M., Saunders, N.F.W., Curmi, P.M., Forwood, J.K., Bodén, M., and Kobe, B. (2011). Molecular basis for specificity of nuclear import and prediction of nuclear localization. Biochimica et Biophysica Acta (BBA) - Molecular Cell Research 1813, 1562–1577. 10.1016/J.BBAMCR.2010.10.013.

50. Nardozzi, J.D., Lott, K., and Cingolani, G. (2010). Phosphorylation meets nuclear import: A review. Cell Communication and Signaling 8, 1–17. 10.1186/1478-811X-8-32/FIGURES/4.

51. Xing, H., Huang, Y., Kunkemoeller, B.H., Dahl, P.J., Muraleetharan, O., Malvankar, N.S., Murrell, M.P., and Kyriakides, T.R. (2022). Dysregulation of TSP2-Rac1-WAVE2 axis in diabetic cells leads to cytoskeletal disorganization, increased cell stiffness, and dysfunction. Scientific Reports 2022 12:1 12, 22474- 10.1038/s41598-022-26337-1.

52. Bermudez, D.M., Herdrich, B.J., Xu, J., Lind, R., Beason, D.P., Mitchell, M.E., Soslowsky, L.J., and Liechty, K.W. (2011). Impaired Biomechanical Properties of Diabetic Skin: Implications in Pathogenesis of Diabetic Wound Complications. Am. J. Pathol. 178, 2215. 10.1016/J.AJPATH.2011.01.015.

53. Nowak, J.A., and Fuchs, E. (2009). Isolation and culture of epithelial stem cells. Methods in Molecular Biology 482. 10.1007/978-1-59745-060-7_14.

54. Rogge, H., Artelt, N., Endlich, N., and Endlich, K. (2017). Automated segmentation and quantification of actin stress fibres undergoing experimentally induced changes. J. Microsc. 268. 10.1111/jmi.12593.

55. Laskowski, R.A., MacArthur, M.W., Moss, D.S., and Thornton, J.M. (1993). PROCHECK: a program to check the stereochemical quality of protein structures. J. Appl. Crystallogr. 26, 283–291. 10.1107/S0021889892009944;CTYPE:STRING:JOURNAL.

56. Colovos, C., and Yeates, T.O. (1993). Verification of protein structures: Patterns of nonbonded atomic interactions. Protein Science 2, 1511–1519. 10.1002/PRO.5560020916;ISSUE:ISSUE:DOI.

57. Eisenberg, D., Lüthy, R., and Bowie, J.U. (1997). [20] VERIFY3D: Assessment of protein models with three-dimensional profiles. Methods Enzymol. 277, 396–404. 10.1016/S0076-6879(97)77022-8.

58. Wiederstein, M., and Sippl, M.J. (2007). ProSA-web: interactive web service for the recognition of errors in three-dimensional structures of proteins. Nucleic Acids Res. 35, W407–W410. 10.1093/NAR/GKM290.

59. Lill, M.A., and Danielson, M.L. (2010). Computer-aided drug design platform using PyMOL. Journal of Computer-Aided Molecular Design 2010 25:1 25, 13–19. 10.1007/S10822-010-9395-8.

60. Warnecke, A., Sandalova, T., Achour, A., and Harris, R.A. (2014). PyTMs: a useful PyMOL plugin for modeling common post-translational modifications. BMC Bioinformatics 2014 15:1 15, 370-. 10.1186/S12859-014-0370-6.

61. Biovia, D.S. (2021). Discovery studio 2021 client, San Diego: Dassault Systèmes. Discovery studio.

