## Supplementary Data for "Actin-dependent mechanotransduction controls nucleocytoplasmic partitioning of DNMT3a through ERK1/2 signaling during cutaneous wound healing"

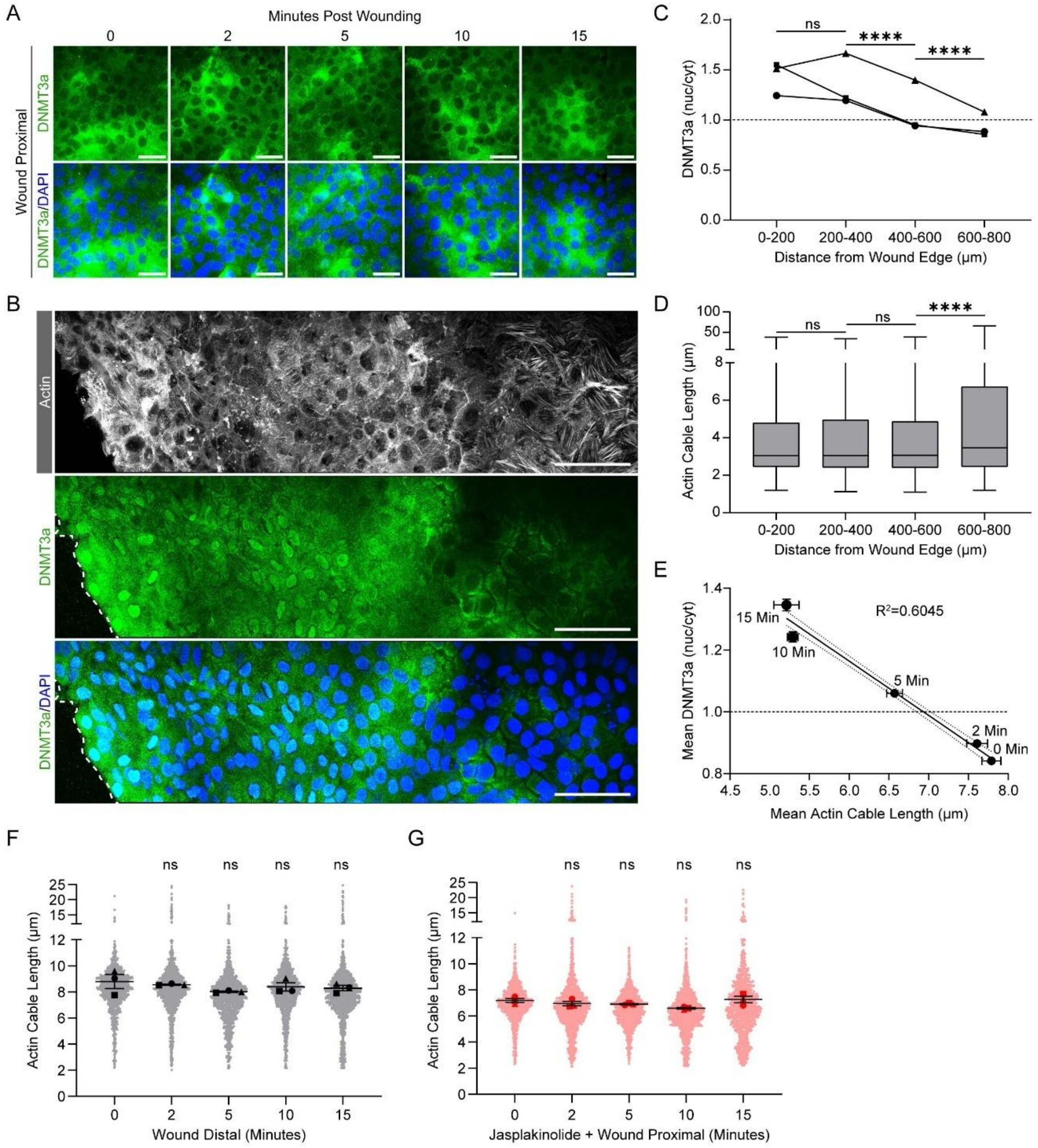


**Figure S1: Spatiotemporal regulation of actin cytoskeleton and nuclear localization of DNMT3a upon wounding.**

**(A)** DNMT3a immunostaining (green) and DAPI (blue) at wound distal (WD) site (> 600 μm from wound edge) showing no change of DNMT3a localization at 5-, 10-, and 15- minutes post wounding. **(B)** Spatial gradient of actin cytoskeleton remodelling (grey) from wound edge (left to right) (Top); Gradient of decreasing nuclear localization (green) from wound edge (left to right) (Middle), DNMT3a and DAPI (blue) merged image (Bottom); captured at 15 minutes post wounding. **(C)** Quantification of DNMT3a (nuc/cyt) from the spatial gradient taking

200*200 µm$^2$ sliced area from the wound edge up to 800 µm showing a decreasing nuclear localization from the wound edge with predominant cytoplasmic localization >600 µm from wound edge. **(D)** Quantification of actin cable length shortening from wound edge. Shorter actin cables up to 600 µm correlates with nuclear localization of DNMT3a. Longer actin cable beyond 600 µm correlates with cytoplasmic localization of DNMT3a. **(E)** Linear regression analysis shows strong correlation between actin cable length shortening and nuclear localization of DNMT3a in a time-dependent manner. The solid line shows a linear fit to the data ($R^2$, squared correlation coefficient). **(F-G)** Quantification of actin cable length in wound distal (WD) and wound proximal (WP) site with Jasplakinolide pre-treatment respectively.

Scale bar: 50 µm for **(A)**, and 100 µm **(B)**. The data are represented as mean ± s.e.m. n=6 for **(A)** and **(E)**; n=3 for (B-D) and **(G)**. P-values in **(C)** and **(D)** were calculated using 2-way ANOVA with Sidak's multiple comparison test where ns= $P > 0.05$, *= $P \leq 0.05$, **= $P \leq 0.01$, ***= $P \leq 0.001$, ****= $P \leq 0.0001$

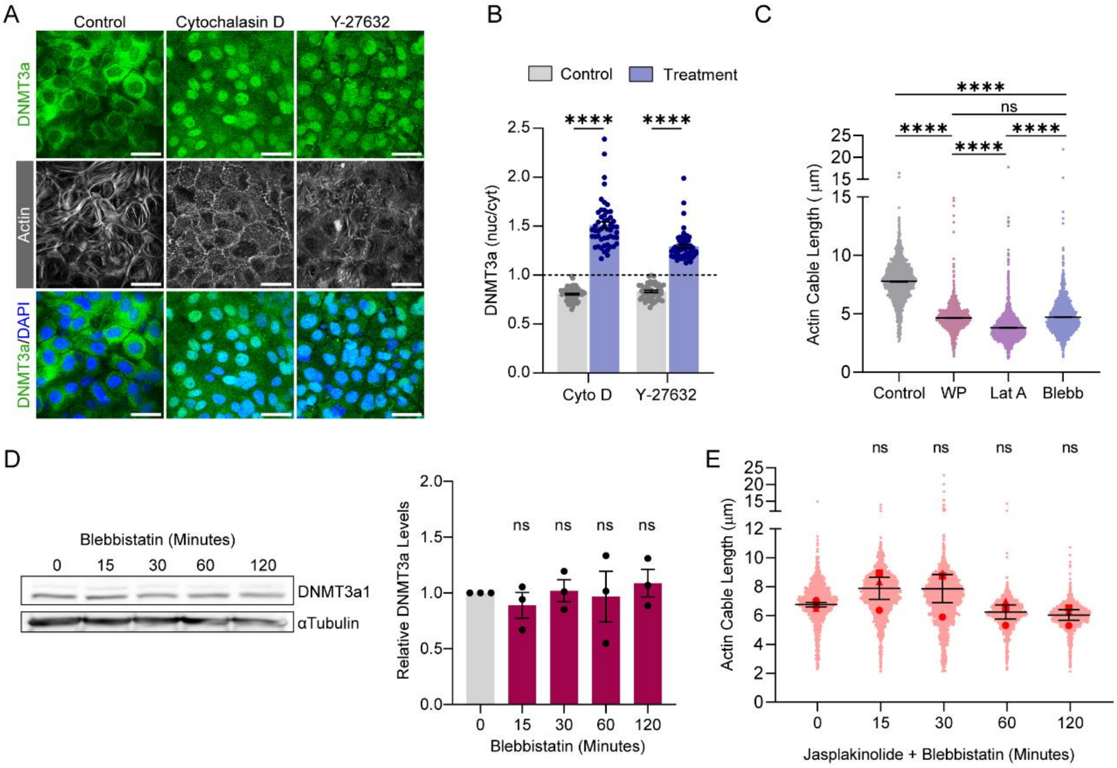


**Figure S2: Partial actin disruption is sufficient for nuclear localization of DNMT3a.**

**(A)** DNMT3a immunostaining (green) with phalloidin (grey), and DAPI (blue) upon Cytochalasin D and ROCK inhibitor, Y-27632 treatment. **(B)** Quantification of DNMT3a (nuc/cyt) upon Cytochalasin D and Y-27632 treatment. **(C)** Quantification and comparison of actin cable length among wound proximal (WP) site, Latrunculin A and Blebbistatin treatment. **(D)** Western blot and quantification of DNMT3a upon Blebbistatin treatment. **(E)** Quantification of actin cable length upon Jasplakinolide + Blebbistatin treatment.

Scale bar: 50 µm. The data are represented as mean ± s.e.m. n=4 for (A-C); n=3 for **(D)**. P-values in **(B-C)** were calculated using 1-way ANOVA with Tukey's multiple comparison test, and (D-E) were calculated using 2-way ANOVA with Sidak's multiple comparison test where ns= P > 0.05, *= P ≤ 0.05, **= P ≤ 0.01, ***= P ≤ 0.001, ****= P ≤ 0.0001.

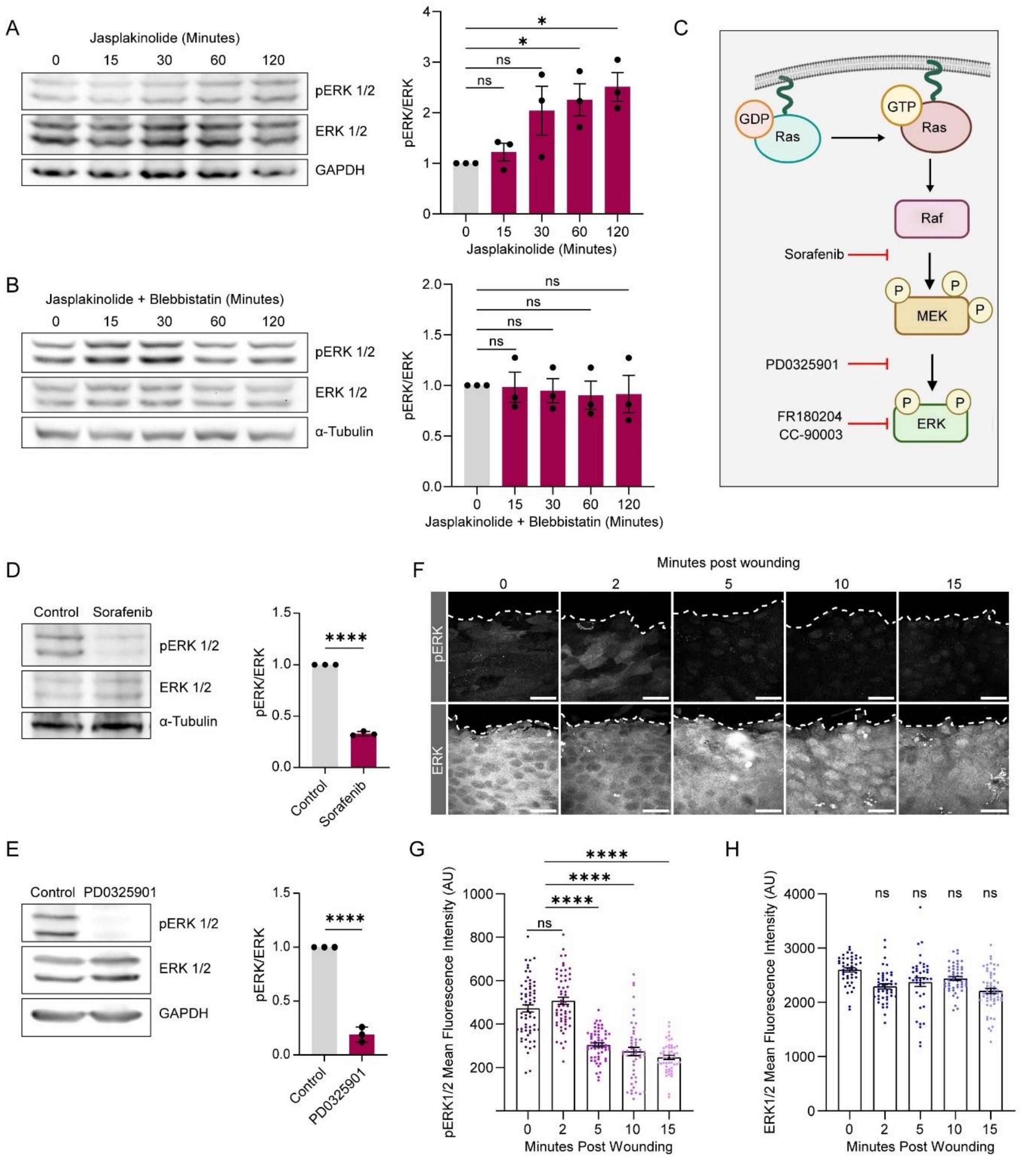


**Figure S3: Actin remodeling-dependent ERK inactivation enables DNMT3a nuclear localization. (A)** Western blot and quantification of pERK/ERK upon Jasplakinolide treatment. **(B)** Western blot and quantification of pERK/ERK upon Jasplakinolide + Blebbistatin treatment. **(C)** Schematic showing targets of ERK signaling pathway inhibitors. **(D-E)** Western blot and quantification of pERK/ERK upon Sorafenib and PD0325901 respectively. **(F)** pERK and ERK immunostaining (grey) at the wound proximal site. **(G-H)** Quantification of Mean Fluorescence Intensity (MFI) of pERK and ERK respectively upon wounding.

Scale bar: 50 μm for **(F)**. The data are represented as mean ± s.e.m. n=3 for (A-H). P-values in **(A)**, **(B)**, **(G)** and **(H)** were calculated 2-way ANOVA with Tukey's multiple comparison test and in **(D)** and **(E)** were calculated using unpaired Welch's t test where ns= $P > 0.05$, *= $P \leq 0.05$, **= $P \leq 0.01$, ***= $P \leq 0.001$, ****= $P \leq 0.0001$.

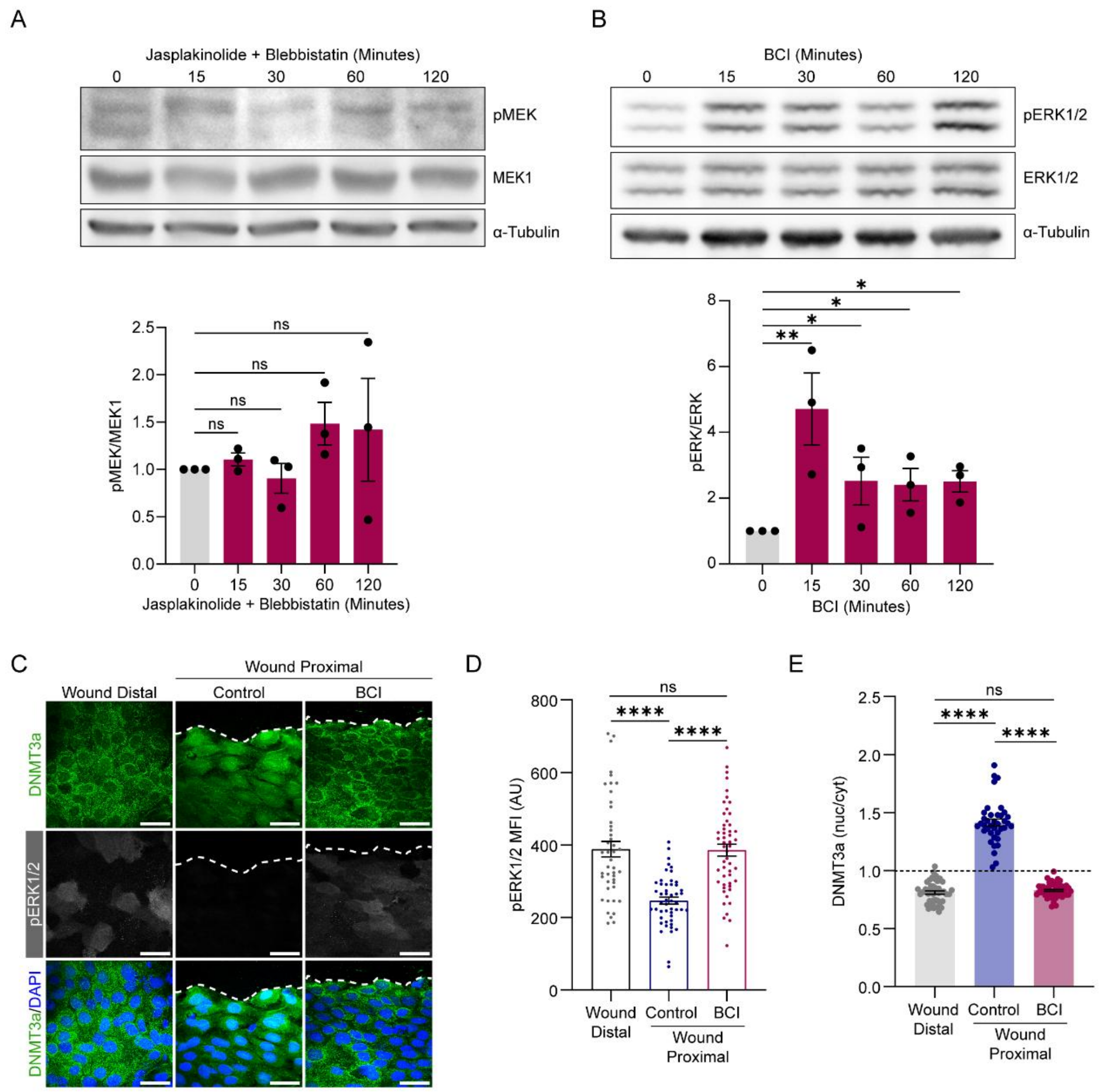


**Figure S4: MEK and DUSP6 coordinate DNMT3a nuclear localization by regulating ERK1/2 inactivation.**

**(A)** Western blot and quantification of pERK/ERK upon Jasplakinolide + Blebbistatin treatment. **(B)** Western blot and quantification of pERK/ERK upon BCI treatment. **(C)** DNMT3a (green) immunostaining with pERK (grey) and DAPI (blue) upon wounding with or without BCI treatment. **(D-E)** Quantification of pERK MFI and DNMT3a (nuc/cyt) respectively upon wounding with or without BCI treatment.

Scale bar: 50 µm for **(C)**. The data are represented as mean ± s.e.m. n=3 for **(A-B)**; n=4 for **(C-E)**. P-values in **(A-B)**, **(D-E)** were calculated using 2-way ANOVA with Tukey's multiple comparison test, where ns= P > 0.05, *= P ≤ 0.05, **= P ≤ 0.01, ***= P ≤ 0.001, ****= P ≤ 0.0001.

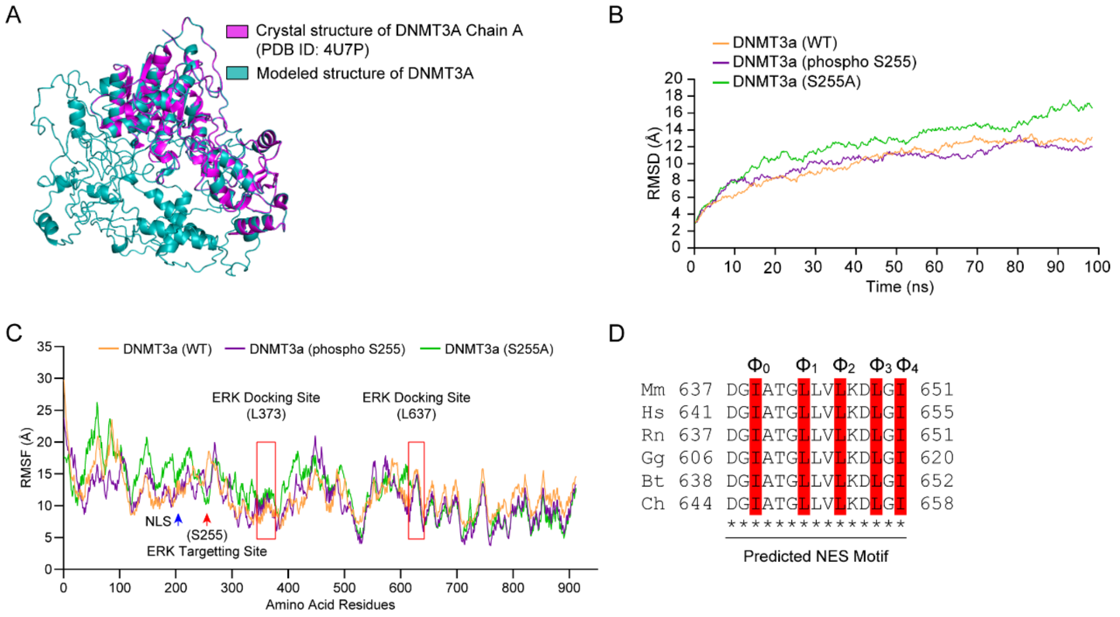


**Figure S5: Phosphorylation at S255 mediates intramolecular masking of the NLS.**

**(A)** Superimposition of modelled DNMT3a structure (cyan) with crystal structure of DNMT3a chain A structure (purple) available on PDB (4U7P). The superimposition RMSD was 0.157 Å **(B)** RMSD analysis of modelled structures of DNMT3a (WT, S255A, and phospho S255). **(C)** Root mean square fluctuation (RMSF) analysis of the modelled structures of DNMT3a (WT, S255A, and phospho S255). ERK docking sites are marked as red rectangles whereas NLS and ERK targeting site (S255) are marked by blue and red arrows respectively. **(D)** Predicted Nuclear Export Signal (NES) sites on DNMT3a with critical hydrophobic residues highlighted in red.

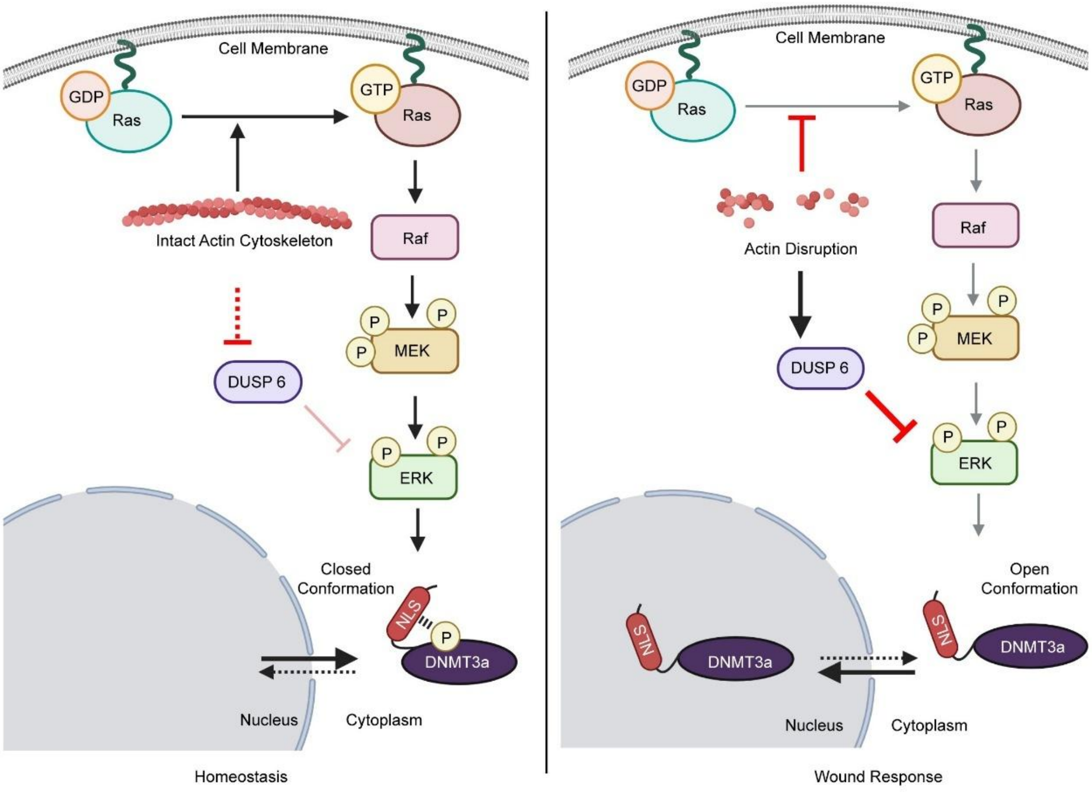


**Figure S6: Schematic of mechanism of regulation of subcellular localization of DNMT3a upon wound-induced mechanical cues**

**Supplementary Table 1:**

| | DNMT3a (wild) | DNMT3a (phospho S255) | DNMT3a S255A |
|---|---|---|---|
| **GLN 248** | 0 | 0.012 | -1.49 |
| **GLN 249** | -0.511 | -2.072 | -5.66 |
| **PRO 250** | 5.86 | 2.96 | 0 |
| **THR 251** | 5.91 | 5.34 | 0 |
| **ASP 252** | -164.7 | -15.45 | 0 |
| **PRO 253** | 1.67 | -3.41 | 0 |
| **ALA 254** | 0.61 | 5.84 | 0 |
| **SER 255** | -8.29 | -225.77 | 0 |
| **PRO 256** | 0.35 | -1.8 | 0 |
| **THR 257** | 1.49 | -1.92 | 0 |
| **VAL 258** | 0 | 0.21 | 0 |
| **ALA 259** | 0 | 0 | 0 |
| **THR 260** | 0 | 0.000023 | 0 |
| **THR 261** | 0 | 0 | 0 |
| **PRO 262** | 0 | 0 | 0 |
| **GLU 263** | 0 | 0 | 0 |

**Supplementary Table 2:**

| | DNMT3a (wild) | DNMT3a (phospho S255) | DNMT3a S255A |
|---|---|---|---|
| **LYS 202** | -123.5 | -235.3 | -4.97 |
| **ARG 203** | -46 | 4.06 | -3.38 |
| **ASP 204** | 14.5 | -4.74 | -0.312 |
| **GLU 205** | 0.19 | 0 | 1.5 |
| **TRP 206** | -2.7 | 0 | 0 |
| **LEU 207** | 0 | 0 | 0 |
| **ALA 208** | 0 | 0 | 0 |
| **ARG 209** | 0 | 0 | 0 |
| **TRP 210** | 0 | 0 | 0 |
| **LYS 211** | 0 | 0 | 0 |
| **ARG 212** | 0 | 0 | 0 |
| **GLU 213** | 0 | 0 | 0 |
| **ALA 214** | 0 | 0 | 0 |
| **GLU 215** | 0 | 0 | 0 |
| **LYS 216** | 0 | 0 | 0 |
| **LYS 217** | 0 | 0 | 0 |
| **ALA 218** | 0 | 0 | 0 |
| **LYS 219** | 0 | 0 | 0 |
